# Pooled single-cell CRISPRa/i screens for functional genomics in bacteria at scale

**DOI:** 10.64898/2025.12.20.695731

**Authors:** Jacob R. Brandner, Quoc Tran, Yujia Huang, Dmitry Sutormin, Karl D. Gaisser, Georg Seelig, Jesse G. Zalatan, James M. Carothers, Anna Kuchina

## Abstract

Pooled CRISPR screens using single-cell RNA sequencing (scRNA-seq) have emerged as powerful tools to uncover gene function, map regulatory networks, and identify genetic interactions. However, the inherent sparsity of bacterial scRNA-seq data has posed a major challenge toward applying these approaches to bacteria. Here, we present mapSPLiT, a pooled bacterial CRISPR activation and interference (CRISPRa/i) screening platform that enables large-scale experiments that simultaneously link hundreds of perturbations to their corresponding single-cell transcriptomes. By targeting 52 known or putative transcription factors with 118 perturbations in a pooled experiment, we expanded the *E. coli* regulatory network map, determined the function of putative regulators, and identified emergent global phenotypes. By targeting combinations of transcription factors simultaneously, we uncovered genetic interactions and regulatory logic between them. Mapping regulatory networks for carbon utilization in *P. putida* revealed control points that could expand metabolic flexibility and improve biomanufacturing. Together, these results establish mapSPLiT as a generalizable platform for single-cell functional genomics in bacteria.

## Introduction

High-content single-cell CRISPR screens in eukaryotic systems provide unprecedented resolution and scale for interrogating gene function and regulatory networks^1–3^. CRISPR-Cas systems use single-guide RNAs (sgRNAs) to programmably target genes of interest, and are readily adapted to massively parallel experiments. By pairing CRISPR-Cas screens to high-dimensional readouts such as single-cell RNA sequencing (scRNA-seq), a single pooled experiment can measure thousands of transcriptome-wide responses to individual perturbations.

Although there are several droplet and combinatorial indexing scRNA-seq strategies in bacterial systems^4–7^, single-cell CRISPR screens have not yet been implemented in bacteria to date. Such screens require simultaneously capturing the transcriptome and a perturbation identifier such as an sgRNA or barcode within the same cell. The significant technical hurdle of sparse transcript capture in bacterial systems, typically tens to hundreds of transcripts per cell, has made it difficult to detect and link CRISPR perturbations to their corresponding cells in large pooled screens^1^.

To overcome these challenges, we developed mapSPLiT (Microbial Analysis of Perturbations using Split-Pool Ligation Transcriptomics) (**Fig. 1A**). mapSPLiT integrates bacterial transcriptional repression (CRISPRi)^8^ and activation (CRISPRa)^9,10^ to perturb gene expression with microSPLiT scRNA-seq technology^4,5^ for transcriptome profiling. To address sparse transcript capture, mapSPLiT employs a separate guide barcode construct (GBC) to identify each sgRNA program in single cells and a PCR-based enrichment protocol to improve GBC detection. To increase the recovery of mRNA reads, we implemented a sequencing-guided rRNA depletion strategy. Together, these innovations enabled large-scale pooled single- and multi-guide single-cell CRISPR screens in *Escherichia coli* (*E. coli*). In one experiment, mapSPLiT scaled to more than a hundred single-guide perturbations and resolved regulatory networks for known and putative transcription factors at single-cell resolution. Additionally, we demonstrated portability across bacterial species by applying mapSPLiT to the non-model microbe *Pseudomonas putida* (*P. putida*) whose regulatory network is largely unknown^11^. Overall, this platform enables scalable single-cell CRISPR screens in multiple bacterial species, advancing fundamental biology and emerging biotechnology.

**Fig. 1.**
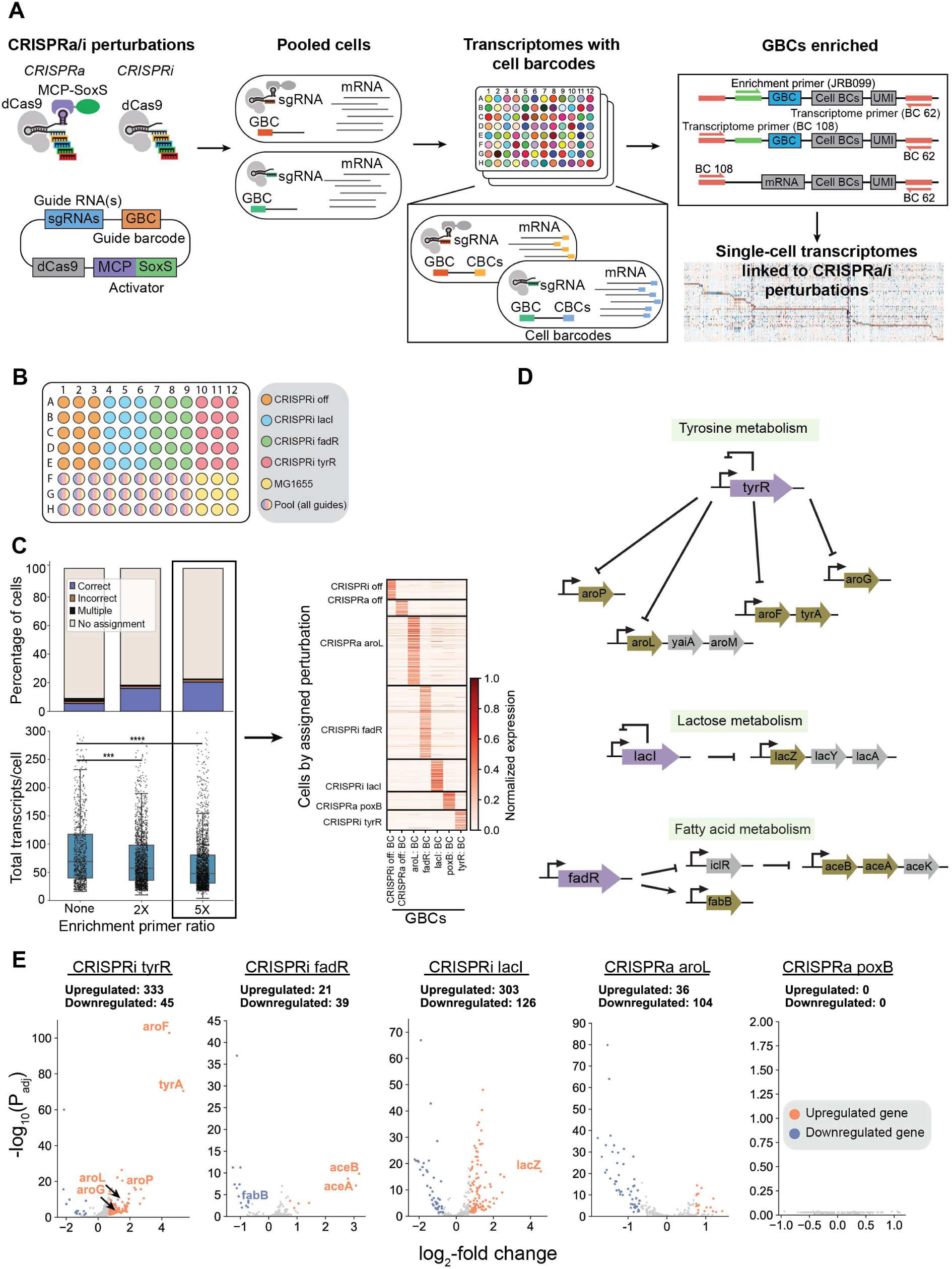
Benchmarking the mapSPLiT platform. **(A)** Experimental overview of mapSPLiT: Microbial Analysis of Perturbations using Split-Pool Ligation Transcriptomics. **(B)** Reverse transcription microSPLiT barcodes for sample identification. **(C)** Top left panel, the percentage of GBC-assigned cells by category (correct, incorrect, multiple, or no assignment) for different concentrations of enrichment primer relative to reverse primer for the arrayed perturbations. Bottom left panel, medians and interquartile ranges (IQRs) of the total transcripts in each cell. Two-sided Welch’s t-test, ***p<0.001. Right panel, normalized expression of all GBC transcripts (columns) for single cells (rows) grouped by GBC-assigned perturbation. **(D)** Depiction of gene regulatory networks found for the CRISPRi-perturbed transcription factors (purple). The DEGs (gold) are organized into operon structures including the non-significant genes (gray). **(E)** Volcano plots highlighting significantly upregulated (orange) and downregulated (blue) genes. Text on the plot indicates that the differentially expressed genes were previously annotated to be within the regulatory network.

## Results

### Guide barcode enrichment enhances assignment of CRISPR programs in single cells

To implement single-cell CRISPR screens in bacteria, we first targeted transcription factors in *E. coli*. Perturbing transcription factors should affect mRNA levels for multiple downstream target genes, producing effects that are detectable even if transcriptome coverage is sparse. Further, because protein abundance in bacteria is primarily determined by the rate of transcriptional initiation^12^, understanding and mapping transcriptional regulatory networks is critical to fully characterizing and predictably engineering bacterial systems. We targeted three transcription factors with CRISPRi repression: tyrosine repressor (*tyrR*)^13^, fatty acid degradation regulator (*fadR*)^14^, and lactose inhibitor (*lacI*)^15^. These transcription factors have highly annotated networks and thus were attractive candidates for our pilot experiment. Additionally, we activated the two enzymes, shikimate kinase II (*aroL)* and pyruvate oxidase (*poxB*), with CRISPRa because these targets were previously validated^10,16^ (**table S1**). We performed perturbations of these five single targets and measured the transcriptional outcomes by microSPLiT.

To implement guide barcodes that specify the CRISPR perturbation at the single cell level, each sgRNA gene was co-expressed with a separate GBC, a 316 bp transcript with a unique 12 bp barcode encoding the perturbation identity at the 5’ end (**Methods**). During the microSPLiT barcoding process, this GBC transcript received the same cellular barcode as every other RNA transcript for that cell, allowing transcript reads to be associated with a GBC and the corresponding CRISPR perturbation. For this initial experiment, instead of delivering a pooled library of sgRNAs, we arrayed both the CRISPR-perturbed and wild-type *E. coli* cells into sample-specific wells in the reverse transcription plate. In this format, the first cell barcode appended during microSPLiT combinatorial barcoding became the sample identifier that specified the corresponding sgRNA perturbation (**Fig. 1B**). This approach allowed us to evaluate whether GBCs would be effective for sgRNA identification in a pooled library format.

To ensure that GBCs were captured in scRNA-seq, we designed a PCR-based enrichment method by adding an additional primer binding site to the 5’ end of the GBC sequence. To prepare the microSPLiT libraries for sequencing, every cDNA transcript was amplified with a universal set of forward and reverse primers. To enrich for the GBC sequence, we introduced the GBC-specific primer at the start of cDNA library amplification (**Fig. 1A and table S2**). The resulting sequencing reads were assigned to single cells based on the microSPLiT cellular barcodes. This process resulted in single-cell transcriptomes with associated GBCs, providing the information necessary to link perturbations to phenotypes.

Ideally, we expect to observe GBCs encoding one perturbation per cell. In practice, for some cells we cannot recover any GBC transcripts, while for others we detect GBCs from multiple perturbations. This could either indicate doublet formation during the scRNA-seq workflow, genuine co-delivery of multiple GBCs to the same cell, or PCR artifacts introduced during GBC enrichment. Thus, for each cell where we recovered GBC transcripts from more than one perturbation, we assigned the cell to the perturbation corresponding to the most abundant GBC (**Methods**). We then independently determined the percentage of correctly and incorrectly assigned cells using the microSPLiT sample identifier as the true identity. Without enrichment, we were able to correctly assign only 5.5% of cells with an incorrect assignment rate of 1.5%. We then quantified how perturbation assignment was affected by GBC enrichment, using multiple enrichment primer concentrations, or by using sgRNA reads themselves. We found that GBC enrichment at 5X primer concentration resulted in the highest rate of correct assignment (21%), 3.8-fold higher than without enrichment, while the rate of incorrect assignment remained at 1.7% (**Fig. 1C, fig. S1, A and B, and note S1**). Therefore, we proceeded with the 5X enrichment GBC condition for all future experiments.

To demonstrate that mapSPLiT can be employed in a pooled screen, we sought to evaluate GBC detection efficiency by delivering all perturbed strains (5 on-targets, 2 controls) together to cells in a single well. As this setup does not allow us to use the microSPLiT sample identifiers, we assigned our single-cell transcriptomes solely with the GBC. In this pooled sample with the 5X primer condition, 25% of cells could be assigned to a perturbation using the GBC, consistent with the assignment rate observed in the arrayed samples (**fig. S1C**). Overall, these results establish that our barcode enrichment strategy is highly accurate, is compatible with pooled screening approaches, and requires minimal changes to the microSPLiT protocol.

### A pooled mapSPLiT experiment uncovers regulatory network responses

With a validated perturbation assignment strategy, we assessed whether a pooled mapSPLiT screen could resolve known transcriptional networks by analyzing *E. coli* transcription factor perturbation responses. We sequenced a library of 44,723 cells from the pilot experiment with an assignment rate of 25% (**fig. S1D**) for a total of 13,046 assigned cells. Assigned cells had a median of 94 transcripts and 78 genes per cell across all CRISPR perturbations (**fig. S2, A to C**). To identify which genes were affected by our perturbations, we applied a Wilcoxon rank sum test on the expression of each gene in comparison to its expression in the off-target controls. For each perturbation we found a set of significantly differentially expressed genes (DEGs) that met our thresholds (p_adj_<0.05, abs(log_2_-fold change) > 0.7, expressed in at least 5% of the perturbation’s cells) (**table S3, Methods**).

To systematically investigate the gene expression responses from the CRISPRi perturbations, we examined the DEGs in each perturbation (**Fig. 1, D and E**). As expected, CRISPRi repression of *lacI* led to upregulation of the *lacZYA* operon. For *fadR* repression, the highest differentially expressed genes were malate synthase (*aceB*) and isocitrate lyase (*aceA*) (**Fig. 1E**), which are not known to be direct targets of *fadR*. However, *fadR* is a direct activator of the repressor *iclR*, which has a known binding site on the *aceBA* promoter^17^. Thus, CRISPRi repression of *fadR* likely decreases expression of iclR, which results in increased expression from the *aceBA* promoter (**Fig. 1, D and E**). Additionally, as expected, we observed downregulation of *fabB*, a known direct target of *fadR* (**Fig. 1, D and E**). For CRISPRi repression of *tyrR*, we observed expected expression of aromatic amino acid biosynthesis genes (**Fig. 1, D and E**). We did not observe significant knockdown of the CRISPRi-targeted regulators themselves (**fig. S2D**), likely due to the sparsity of single-cell data and the inherent low expression of transcription factors^18^. Nonetheless, we demonstrated that mapSPLiT is capable of uncovering downstream expression changes matching known regulatory responses.

In the CRISPRa poxB strain, we did not detect significant expression changes for poxB or other genes. We used RT-qPCR to confirm that poxB CRISPRa is effective, so the lack of a detectable effect in mapSPLiT suggests that the increased poxB levels and any associated downstream effects remain below the limit of detection in this experiment (**Fig. 1E, fig. S2D, fig. S3, note S2**).

For the *aroL* CRISPRa perturbation, we expected to increase metabolic flux through the aromatic amino acid biosynthesis pathway. Accordingly, we observed upregulation of the phenylalanine and tyrosine tRNA synthetase genes (*pheT*, *pheS*), which could indicate a surplus of these amino acids. We also observed transcriptional changes in several other glycolysis genes, consistent with prior data from another aromatic acid overproduction strain (L-phenylalanine)^19^ (**fig. S4, B to D, note S2**). We did not directly observe activation of the *aroL* gene itself (**fig. S2D**). One potential explanation is that *aroL* was initially upregulated, which then had downstream effects that were detectable, but the *aroL* mRNA levels were no longer elevated at the time of measurement. Altogether, the mapSPLiT platform is capable of measuring CRISPRa and CRISPRi regulatory network responses. These results, and our success with GBC enrichment, provided a foundation for scaling our platform to multiple guides and larger pooled screens.

### rRNA depletion improves transcript capture for mapSPLiT

In initial experiments, we observed that a large fraction of sequencing reads were ribosomal RNA (rRNA). rRNA comprises over 95% of all transcripts in *E. coli*, but rRNA reads generally provide little information about transcriptional responses and are typically removed from analysis following sequencing^20^ thus incurring unnecessary sequencing costs. Methods to remove rRNA biochemically prior to sequencing exist^21,22^. Our initial experiments showed that rRNA coverage was non-uniform (**Fig. 2A**), likely because microSPLiT capture and cDNA conversion happens *in situ* with ribosomal and other proteins that may block reverse transcription. To address this non-uniform coverage, we developed a sequencing-guided Cas9 nuclease-based rRNA depletion method to preferentially remove regions of the rRNA sequence that were overrepresented in mapSPLiT sequencing. We selected a subset of 253 sgRNAs, from 824 possible sgRNAs mapping to rRNA sequences, that targeted these overrepresented regions (**Fig. 2A, Methods**). The purified cDNA library obtained from the multi-guide mapSPLiT experiment (see next section) was incubated with recombinant Cas9 and the sequencing-guided sgRNA pool to cleave rRNA cDNA. We separately sequenced this depleted library in parallel with an undepleted cDNA library. With rRNA depletion, the mRNA fraction increased from 4.6% to 60.2% (**Fig. 2B**) and median mRNA transcripts per cell more than doubled, from 54 to 115 (**Fig. 2C**), without changing the overall mRNA expression profile (**Fig. 2D, note S3**). The sequencing-guided rRNA depletion method is a cost-effective solution allowing mapSPLiT to scale to larger perturbation libraries by improving detection of transcriptional responses. We used this rRNA depletion method for all subsequent mapSPLiT experiments.

**Fig. 2.**
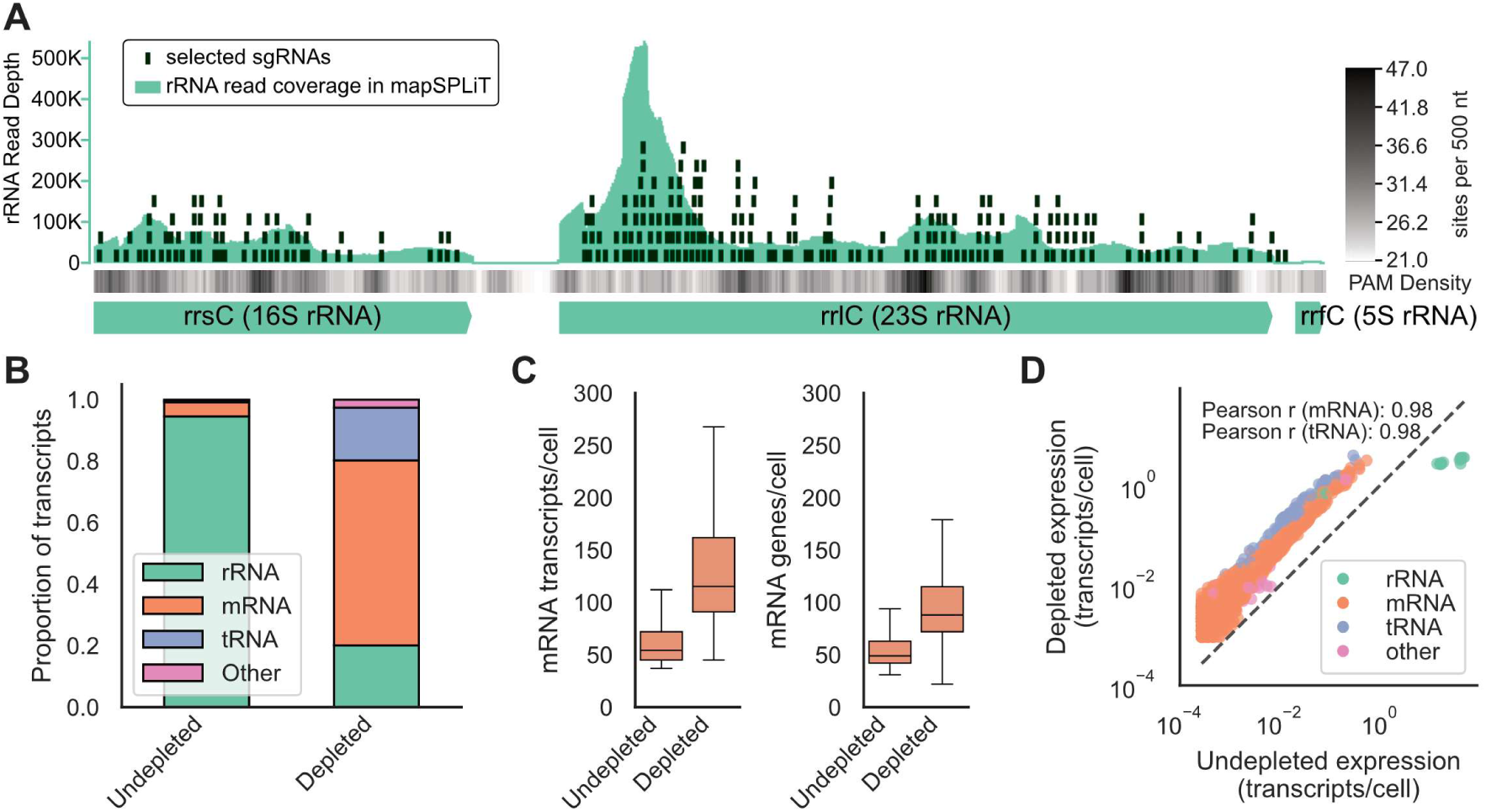
Sequencing-guided rRNA depletion. **(A)** A library of 253 sgRNAs for Cas9 cleavage (black rectangles) was designed to fully cover *E. coli* rRNA sequences, guided by both the local density of PAM sites (grayscale intensity bar) and sequencing depth of rRNA reads (green peaks). The library of cleavage sgRNAs is shown aligned to a representative rRNA gene (rrnC locus). **(B)** Relative proportion of RNA transcripts between samples with and without sequencing-guided rRNA depletion calculated after genome alignment but before single cell filtering. **(C)** Median and IQRs of mRNA transcripts and genes per cell with and without sequencing-guided rRNA depletion calculated after cell filtering with identical parameters. **(D)** Each unique gene was plotted with its expression in the with (y-axis) and without (x-axis) sequencing-guided rRNA depletion calculated as its number of transcripts in each library divided by the total number of filtered cells. The dashed line shows identical gene expression between both conditions.

### Multi-guide mapSPLiT uncovers transcriptome-wide interactions and transcription factor logic

We next evaluated the effects of multiple simultaneous CRISPR perturbations. Non-additive genetic interactions arise when the phenotype of a multi-gene perturbation deviates from that predicted from single-gene effects. As one illustration, nearly half of regulated promoters in *E. coli* are controlled by more than one transcription factor^23^, and their combined activities can produce synergistic^24^ or buffering^25^ outcomes. These interactions occur for co-regulated genes and can extend to downstream genes across the transcriptome.

To probe multi-gene interactions, we designed a pooled multi-guide mapSPLiT experiment. Two or three genes were targeted simultaneously by expressing multiple sgRNAs from a plasmid containing a single GBC (**Fig. 3A, tables S1 and S2**). We focused on the aromatic amino acid biosynthesis pathway and the lactose degradation pathway. In the same cell, we activated a metabolic gene (*aroF*, *aroH, aroL*, *lacZ*) with CRISPRa and repressed its annotated transcriptional regulators (*tyrR* and *trpR* for *aroF*, *aroH*, and *aroL*; *lacI* for *lacZ*) with CRISPRi. This design was expected to increase metabolic pathway activity by simultaneously driving metabolic gene expression and relieving transcriptional repression, thereby exposing synergistic or buffering interactions. For each metabolic target, we tested all combinations of perturbations involving the metabolic gene itself and its regulators (ten two-guide cassettes and eight three-guide cassettes), yielding 18 multi-guide perturbations, including off-target controls (**table S2**). Each strain was grown separately and analyzed in a pooled mapSPLiT experiment, which captured a median of 137 transcripts and 99 genes per cell across 122,763 cells (**fig. S5**).

**Fig. 3.**
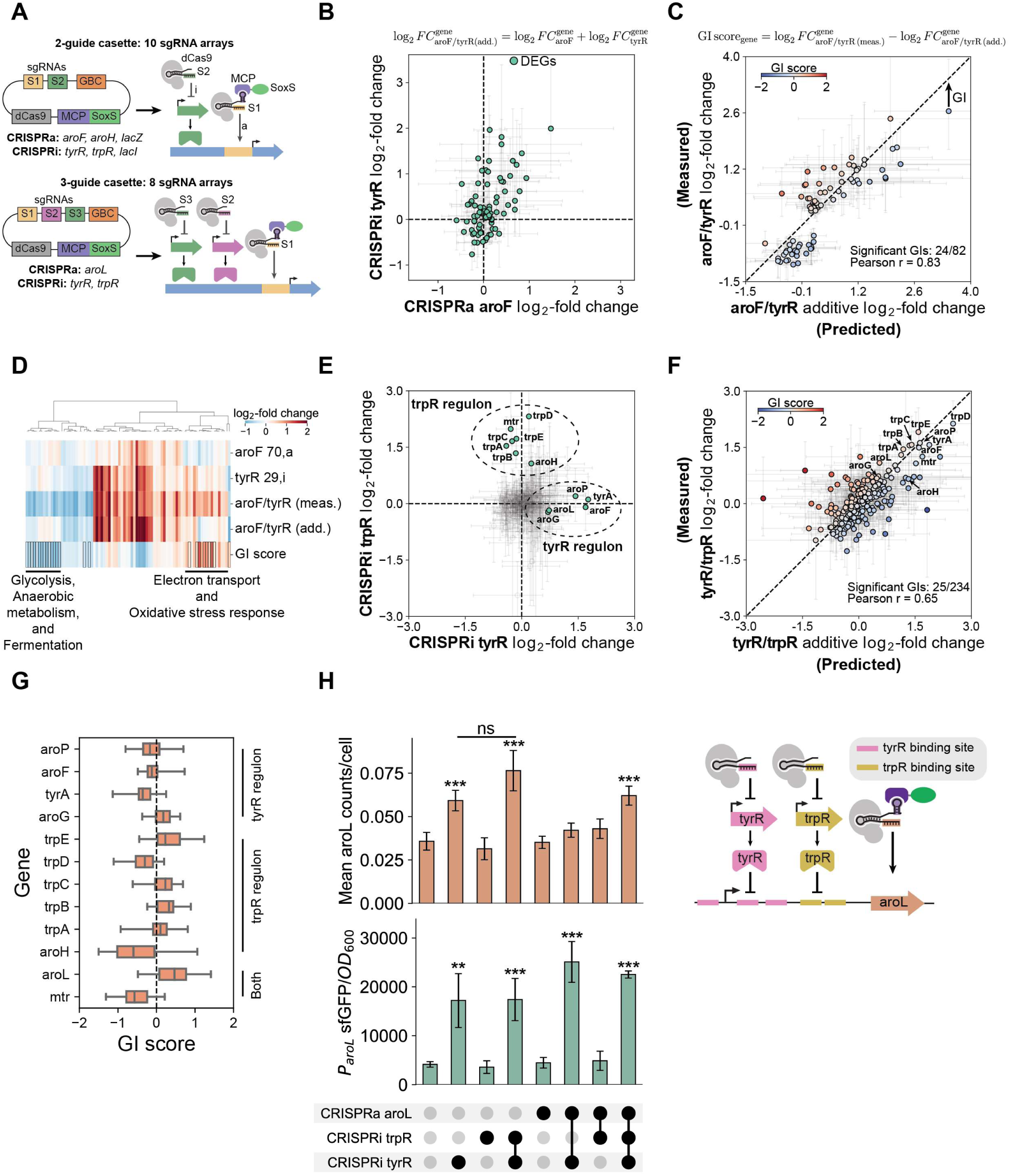
Multi-guide CRISPR screen reveals the dominant regulators at the *aroL* and *mtr* promoters. **(A)** Schematic depicting multi-guide cassettes and targeted genes. “a” indicates activation and “i” indicates inhibition. **(B and E),** Single sgRNA log_2_-fold changes of DEGs for CRISPRa of *aroF* and CRISPRi of *tyrR* **(B)** or the CRISPRi of *tyrR* and CRISPRi of *trpR* **(E)** perturbations relative to an off-target control. Error bars are bootstrapped 95% confidence intervals (CIs). Green dots and text indicate CRISPRa target gene or regulon of targeted transcription factor. **(C and F)** Genetic interaction map for CRISPRa of *aroF* and CRISPRi of *tyrR* **(C)** or CRISPRi of *tyrR* and CRISPRi of *trpR* **(F)** depicting the measured (x-axis) and predicted additive (y-axis) log_2_-fold change for each DEG relative to an off-target control. Error bars are as in (**B,E**). Colorbar shows the genetic interaction (GI) score as the difference between the measured and predicted log_2_-fold changes. Pearson correlations are computed using the mean log_2_-fold changes. Text indicates CRISPRa target gene or regulon of targeted transcription factor. **(D)** log_2_-fold changes of single-guide, multi-guide (“meas.”), and predicted expression (“add.”), alongside GI scores for all 82 DEG s of *aroF* CRISPRa and *tyrR* CRISPRi. The log_2_-fold changes of expression were calculated relative to an off-target control. Black outlines mark significant GI scores (bootstrapped 95% CIs that do not include zero). Columns ordered by hierarchical clustering. **(G)** GI scores for CRISPRi of *tyrR* and CRISPRi of *trpR* for the solely *tyrR* regulated, solely *trpR* regulated, and the co-regulated genes. Whiskers show the boostrapped 95% CI. **(H)** Top left, mean *aroL* transcripts per cell. Bottom left, mean normalized fluorescence (sfGFP/OD_600_) from a P_aroL_ plasmid reporter. Empirical p-values (transcriptomics) or two-sided Welch’s t-test (fluorescence), *p<0.05, **p<0.01, ***p<0.001 (**Methods**). Reporter error bars: SD of 3 strains across two experiments. Black circles indicate contributing sets. Right, schematic of CRISPRa targeting *aroL* and CRISPRi targeting *tyrR* and *trpR* at the *aroL* promoter.

We quantified transcriptome-wide genetic interactions across the metabolic gene/transcriptional regulator pairs. We defined a genetic interaction score for each gene as its measured expression under dual-perturbation minus its predicted additive expression, calculated by summing the effects of each single-gene perturbation (**Methods**). Additive predictions captured most responses (789/1025 genetic interactions). Among the remaining 236 genetic interactions, 101 showed higher-than-predicted changes in gene expression (synergistic) and 135 showed lower-than-predicted changes (buffering) (**fig. S6 and S7**). This framework enables a quantitative assessment of how combinations of genes shape transcriptional network responses.

To examine these interactions in greater depth, we focused on genetic interactions produced by CRISPRa of *aroF* paired with CRISPRi of *tyrR* (*aroF*/*tyrR*) (**Fig. 3, B and C**). This combination produced the largest number of regulated genes with non-additive genetic interactions among the two-guide CRISPRa and CRISPRi perturbations (**tables S4 and S5**). The *aroF/tyrR* perturbation revealed coordinated buffering of glycolysis, fermentation, and anaerobic metabolism genes alongside synergistic activation of electron transport and oxidative stress pathways, indicating broad metabolic rewiring linked to aromatic amino acid biosynthesis (**Fig. 3D, note S4**). These results show that combined perturbations of metabolic and regulatory genes can generate pathway-level genetic interactions and uncover features of network organization.

As a case study in transcription factor logic, we analyzed interactions between *tyrR* and *trpR* at their co-regulated genes *aroL* and *mtr*. Single-gene perturbations confirmed the expected regulon responses. CRISPRi of *tyrR* increased expression of *tyrR*-regulated genes, and CRISPRi of *trpR* increased expression of *trpR*-regulated genes. (**Fig. 3E, fig. S8A**). At the co-regulated genes, *aroL* responded primarily to *tyrR*, whereas *mtr* responded primarily to *trpR* (**Fig. 3, E, G and H**), consistent with prior evidence for dominant regulators at these loci^26–28^. Dual perturbations of *tyrR* and *trpR* at the same time produced no statistically significant, non-additive effects on *aroL* or *mtr* (**Fig. 3, F and G**). Despite prior expectations that both *tyrR* and *trpR* contribute to *aroL* expression^22,23^, fluorescent reporter and mapSPLiT data indicated the *tyrR* alone regulates *aroL* (**Fig. 3H, note S5**). Direct CRISPRa of *aroL* did not further increase expression beyond relief of *tyrR* repression, implying that the promoter may be already maximally activated (**Fig. 3H**). At *mtr*, *tyrR* functioned as an activator and *trpR* as a repressor, yielding an additive response in which repression prevailed (**fig. S8B, note S5**), consistent with prior findings^28^. These findings clarify two distinct modes of promoter regulation. *aroL* is governed solely by *tyrR*, while *mtr* integrates opposing inputs from *tyrR* and *trpR* with repression dominating. This distinction demonstrates how promoters integrate inputs from multiple regulators and preferentially drive transcriptional outcomes.

### Scaled single-cell CRISPR screens uncover novel and known bacterial regulatory networks

We evaluated whether mapSPLiT could scale to over 100 transcription factor perturbations in a high-throughput pooled experiment. We targeted 52 genes with 118 single-perturbation CRISPR sgRNAs (36 CRISPRa, 82 CRISPRi) (**tables S1 and S2**). Of these 52 genes, 51 were regulators, and 24 regulators were “well-characterized” while 27 regulators were “partially characterized” according to the EcoCyc database^29^ (**Methods**). The “well-characterized” targets spanned both global and local regulators with diverse roles across the genome. The “partially characterized” targets included regulators with experimental characterization consisting of transcription factor binding data, perturbation expression data (such as knockouts, knockdowns, or targeted overexpression), or none at all. Our general strategy was to upregulate activators with CRISPRa and downregulate repressors with CRISPRi to increase downstream gene expression and generate detectable regulatory responses. For “partially characterized” regulators, we designed both CRISPRa and CRISPRi sgRNAs where possible (**Methods**).

We performed mapSPLiT and obtained high-quality data sufficient to detect regulatory changes. We sequenced 365,000 cells and retained 76,068 of them after assignment, with a median of 325 transcripts and 161 genes per cell. These data include 104 of 118 perturbations with sufficient resolution, which we defined as at least 100 cells following standard approaches in scRNA-seq perturbation screens^3,30^ (**Fig. 4A**, fig. S9, S10, A and B).

**Fig. 4.**
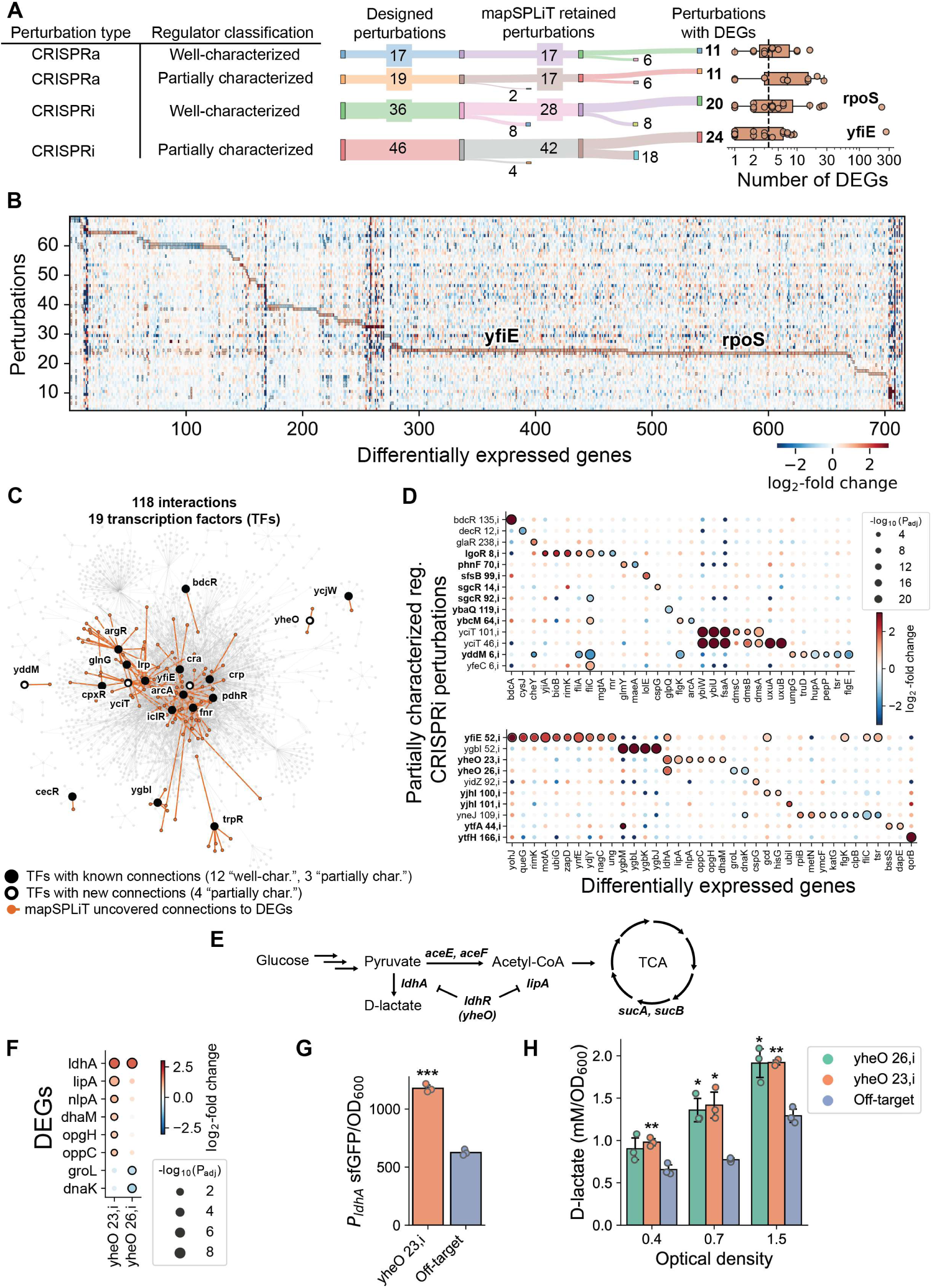
Single-cell CRISPR screens map known and putative regulatory networks for a library of perturbations. **(A)** Left, Sankey diagrams for each perturbation type show numbers of sgRNAs that were designed and ordered, retained during mapSPLiT (at least 100 cells/perturbation after cloning, sequencing, and post-processing), then had at least one DEG (**Methods**). Right, Median and IQRs of the number of DEGs in each retained perturbation (dots), organized by perturbation type. The dotted line indicates the total median number of DEGs (3.5). CRISPRi of *rpoS* and CRISPRi of *yfiE* are labeled because they had over 100 DEGs. **(B)** Expression (relative to an off-target) of DEGs in all perturbations (columns) across all retained perturbations (rows). Black outlines indicate significance (**Methods**). **(C)** Global *E. coli* network of transcription factors that mapSPLiT recapitulated known connections for (closed black nodes), that mapSPLiT found new connections for (open black nodes), or that were unperturbed (gray nodes). Nodes are shown with previously annotated regulatory connections (gray edges) or those found in mapSPLiT (orange edges). **(D)** Expression of up to 10 DEGs for each perturbation (columns) for each “partially characterized” perturbation split between two plots. Bolded perturbation names target transcription factors with no prior expression data. The dot sizes represent the adjusted p-values from the Wilcoxon rank sum test relative to an off-target control. Outlines indicate statistical significance. **(E)** *E. coli* metabolic pathway of D-lactate production. **(F)** Expression of all DEGs (columns) for both *yheO* perturbations (rows). The size and outlines of the dots are as in (**E**). Both *yheO* perturbations produced similar effects with different levels of statistical significance. **(G)** Mean normalized fluorescence (sfGFP/OD_600_) for *yheO* perturbation strains transformed with a P_ldhA_ plasmid reporter. Error bars are standard deviations of three strains. Two-sided Welch’s t-test, ***p<0.001. **(H)** Mean D-lactate production at different *E. coli* growth stages (relative OD_600_ values of 0.4, 0.7, and 1.5) for both *yheO* perturbations. Error bars and statistical significance are as in (**G**).

After stringent filtering for perturbations that had at least one DEG (**Methods**), we retained 66 perturbations covering 43 unique gene targets (**Fig. 4A**). Of these gene targets, 20 were “well-characterized” and 23 were “partially characterized”. Across all 66 perturbations, we found 717 unique DEGs (**Fig. 4B, table S6**). The median number of DEGs per perturbation was 3.5, but some targets resulted in hundreds of DEGs, including the sigma factor *rpoS* and the “partially characterized” *yfiE* gene (**Fig. 4, A and B**). A subset of our CRISPRa perturbations did not activate and sometimes repressed their target genes (**fig. S10C**). There is precedent for CRISPRa complexes having position-dependent repressive effects both in bacterial^31^ and eukaryotic systems^32^.

We first benchmarked regulators against their known regulons by performing a gene regulatory network (GRN) analysis. Using the differential gene expression profiles for each perturbation (**Fig. 4B**), we distinguished the direct and downstream regulatory effects using transcription factor binding and expression data^29,33^ (**Methods**). This analysis identified 100 known direct regulatory network connections for 15 transcription factors with previously-known connections (12 “well-characterized” and 3 “partially characterized”) (**Fig. 4C, fig. S11A**). Generally, we found that perturbations targeting the same regulator produced similar regulatory network responses (**fig. S11A**). 94 of the 100 network connections were unique to a single transcription factor, consistent with the idea that most regulated *E. coli* genes are controlled by two or fewer regulators^23^.

Given that mapSPLiT could recapitulate GRN connections, we then analyzed the gene expression data for our “partially characterized” regulators, finding 4 transcription factors with new connections (*yheO*, *yddM*, *yfiE*, *yciT*) (**Fig. 4C**). These regulators all had prior binding site data but no perturbation expression data for some or all of their putatively annotated promoters. We found DEGs consistent with the putative binding sites, thus, expanding the known *E. coli* GRN with 8 new direct transcriptional regulatory connections (**Fig. 4C, fig. S11, note S6**).

We then used mapSPLiT to infer biological functions for 12 “partially characterized” regulators (*ytfA*, *ytfH*, *yjhI*, *yddM*, *yfiE*, *phnF*, *lgoR*, *sgcR*, *sfsB*, *ybcM*, *yheO*, *ybaQ*), all of which have no prior perturbation expression data. These include a pseudogene (*ytfA*) which we considered “partially characterized” in our analyses. Of these 12 perturbations, CRISPRi of *ytfA* affected expression of a cell division gene (*dapE*) and a biofilm formation regulator (*bssS*). CRISPRi of *yfiE* affected over 300 genes, including ones in the glycerol degradation pathway enzymes (*gldA*, *dhaL*, *dhaK*). Across all these “partially characterized” regulators, we uncovered functions ranging from chemotaxis to stress adaptation, reflecting the diversity of transcription factor roles uncovered by mapSPLiT (**Fig. 4D, fig. S11B, note S6**).

We further experimentally characterized the “partially characterized” regulator *yheO*, prompted by mapSPLiT transcriptional responses that suggested a role for *yheO* in controlling pyruvate flux (**Fig. 4E**). Across two CRISPRi perturbations, mapSPLiT transcriptional responses suggested that *yheO* regulates *ldhA* and *lipA* (**Fig. 4F**). To independently validate our transcriptomic data, we confirmed that *yheO* represses the *ldhA* promoter using a fluorescent reporter assay (**Fig. 4G**). Because *ldhA* encodes a D-lactate dehydrogenase that converts pyruvate to D-lactate, we hypothesized that *yheO* functions as a regulator of D-lactate metabolism. We evaluated this model and observed that repression of *yheO* resulted in significantly higher D-lactate levels (**Fig. 4H**). These transcriptomic and metabolic data support renaming *yheO* as the lactate dehydrogenase repressor (*ldhR*) and links its regulation of D-lactate to pyruvate flux (**note S7**). This data-driven progression, from mapSPLiT data, to validated transcriptional profiles, to metabolite outputs, demonstrates that mapSPLiT generates testable hypotheses for mapping bacterial regulatory networks at scale.

### Single-cell clustering reveals convergence and divergence of perturbation phenotypes

Beyond individual regulators, we compared global transcriptional states to identify distinct and shared responses between perturbations. To investigate these emergent global states, we combined cells for all perturbations of the same transcription factor and clustered cells based on DEGs from all perturbations using Uniform Manifold Approximation and Projection (UMAP)^34^. This analysis revealed 11 clusters corresponding to distinct transcriptional states, annotated by top marker genes. We linked transcriptional states to perturbations by identifying perturbations with the largest fraction of cells in each cluster (**Fig. 5A, fig. S13A, table S7, Methods**). We observed that 4 of the 11 transcriptional clusters were exclusively formed by single perturbations. In addition, we uncovered two additional modes of regulatory behavior: 1) divergence, where a single perturbation resulted in multiple, distinct transcriptional states, and 2) convergence, where a single transcriptional state results from multiple, distinct perturbations (**Fig. 5B, fig. S13B**).

**Fig. 5.**
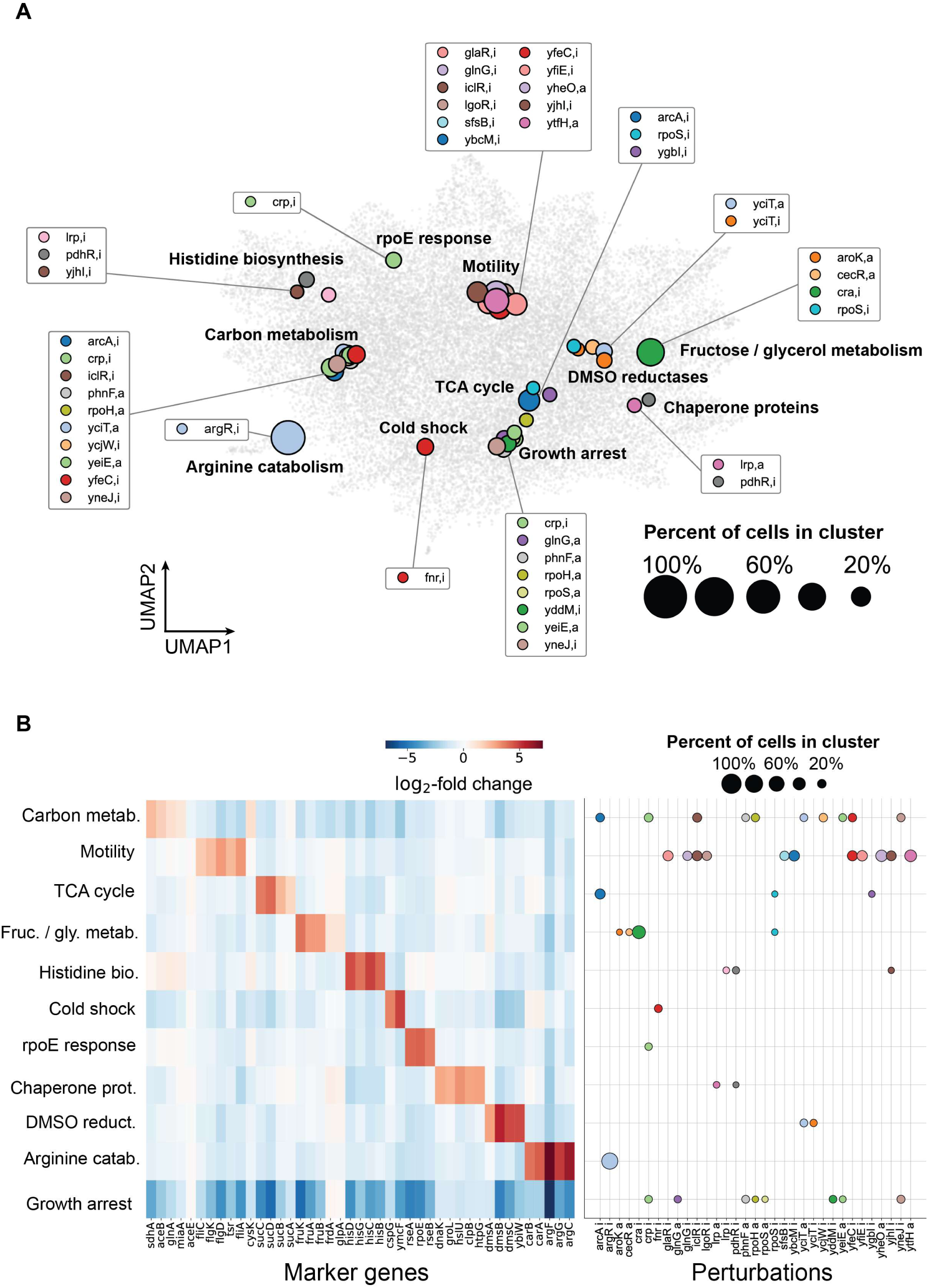
Single-cell clustering reveals perturbation-driven global cell states. **(A)** UMAP clustering where each gray dot represents a single cell. The location of a colored dot represents the average location of a CRISPR perturbation within a cluster; its size represents the proportion of cells in that cluster. Clusters were manually annotated using the marker gene expression profile. **(B)** Left, expression (log_2_-fold change between cells in and out of the cluster) of up to the 5 top unique marker genes, by the absolute z-score from a Wilcoxon rank sum test, (columns) in each cluster (rows). Right, colored dots label a CRISPR perturbation (rows) within the same clusters as the heatmap (rows). The dot sizes represent the proportion of cells assigned to a perturbation within each cluster, and its color is identical to its color on the UMAP.

Single perturbation clusters may indicate direct regulatory control, as around half of regulated promoters in *E. coli* are controlled by a single transcription factor^23^. For 3 of the 4 single perturbation clusters, marker genes for the clusters matched their respective regulons. The fourth cluster resulted from knockdown of *fnr,* which regulates the transition from aerobic to anaerobic growth^35^. We uncovered that Fnr regulates additional cold shock genes (**fig. S14**), which have not been reported as Fnr targets^35^.

We identified 12 divergent perturbations that were each dispersed across multiple transcriptional states (**Fig. 5, fig. S13**). Distinct transcriptional responses, indicative of divergence, have been previously identified for stress and nutrient response regulators^36^. In our dataset, cells with perturbation of the global *crp* transcription factor, which regulates over 180 promoters were distributed across 3 separate clusters: carbon metabolism, rpoE response, and growth arrest^37^. 11 other perturbations were present in 2 clusters each, suggesting that these regulators also had broad transcriptional responses. 5 of these were “well-characterized” regulators (*arcA*, *iclR*, *pdhR*, *rpoH*, *rpoS*) that have been annotated to have large impacts on *E*. *coli* physiology^38–42^. The remaining 6 “partially characterized” perturbations (*phnF*, *yciT*, *yeiE*, *yfeC*, *yjhI*, *yneJ*) were overrepresented in the carbon metabolism, motility, growth arrest, and DMSO reductase clusters, suggesting that they may play roles in the global regulatory network.

We observed 7 of the 11 total transcriptional states were composed of multiple perturbations (**Fig. 5, fig. S13**), indicative of convergence. In *E. coli*, many transcription factors elicit similar gene expression responses^43^. In our dataset, the motility cluster was composed of 11 co-clustered perturbations, consistent with the idea that many different nutrient conditions and stress cues cause changes in flagellum and chemotaxis gene expression^44^. The histidine biosynthesis cluster contained 3 co-clustered perturbations (*lrp*, *pdhR*, *yjhI*), none of which was previously shown to regulate expression of the histidine genes^45^.

We further examined the histidine biosynthesis cluster as a case study of regulatory convergence (**Fig. 5, fig. S13, S15A**). Lrp regulates around a third of the *E. coli* genome via direct and indirect regulatory connections including genes in the isoleucine biosynthesis pathway^46^. Although Lrp has been implicated in responding to histidine^47^, this is the first indication that Lrp regulates histidine biosynthesis. Downregulation of the pyruvate utilization regulator, *pdhR*^40^, appeared to produce a compensatory response that could increase the concentration of NADH, which is a product of histidine biosynthesis (**note S8, fig. S15, B to D**). Inclusion of *yjhI* within the histidine biosynthesis cluster hints at a possible regulatory function for this “partially characterized” transcription factor. Altogether, these results highlight that the single-cell resolution of mapSPLiT can map higher-order associations between individual transcriptional programs governed by each regulator.

### mapSPLiT can be generalized to the industrially-relevant bacterium P. putida

We applied mapSPLiT to uncover regulatory networks in *P. putida*^48^, an emerging chassis in industrial microbiology whose regulatory architecture remains relatively uncharacterized^11^. Specifically, we performed mapSPLiT with CRISPRi targeting three regulators: catabolite repressor activator (*cra*, *PP_0792*), lactate dehydrogenase regulator (*pdhR*, *PP_4734*), and glyoxylate carboligase (*gclR*, *PP_4283*). Each of these regulators has some experimental evidence supporting its known regulon^49–53^, though the extent of characterization varies. Alongside these perturbations of regulators involved in distinct carbon utilization pathways, we also benchmarked whether mapSPLiT could detect CRISPRa responses in *P. putida* by targeting a heterologous fluorescent reporter^54^.

We first assessed whether the mapSPLiT protocol could be ported to *P. putida* without modification by performing the microSPLiT, GBC enrichment, and rRNA depletion workflow as developed for *E. coli* (**Fig. 6A**). Using only the GBC, we achieved 44.6% correct perturbation assignment (**Fig. 6B**), a 2.1-fold improvement over *E. coli*. Using both GBC and sgRNA reads further improved correct assignment to 57.4% (25.0% correct assignment with sgRNA alone) (**Fig. 6B, note S9**). We used both GBC and sgRNA reads for assignment for all downstream analyses. With our sequencing-guided rRNA depletion protocol, the mRNA fraction increased from 2.2% to 10.4% leading to the median capture of 85 transcripts per cell (**fig. S16**), indicating that all of our biochemical workflows were transferable to *P. putida* (**note S9**).

**Fig. 6.**
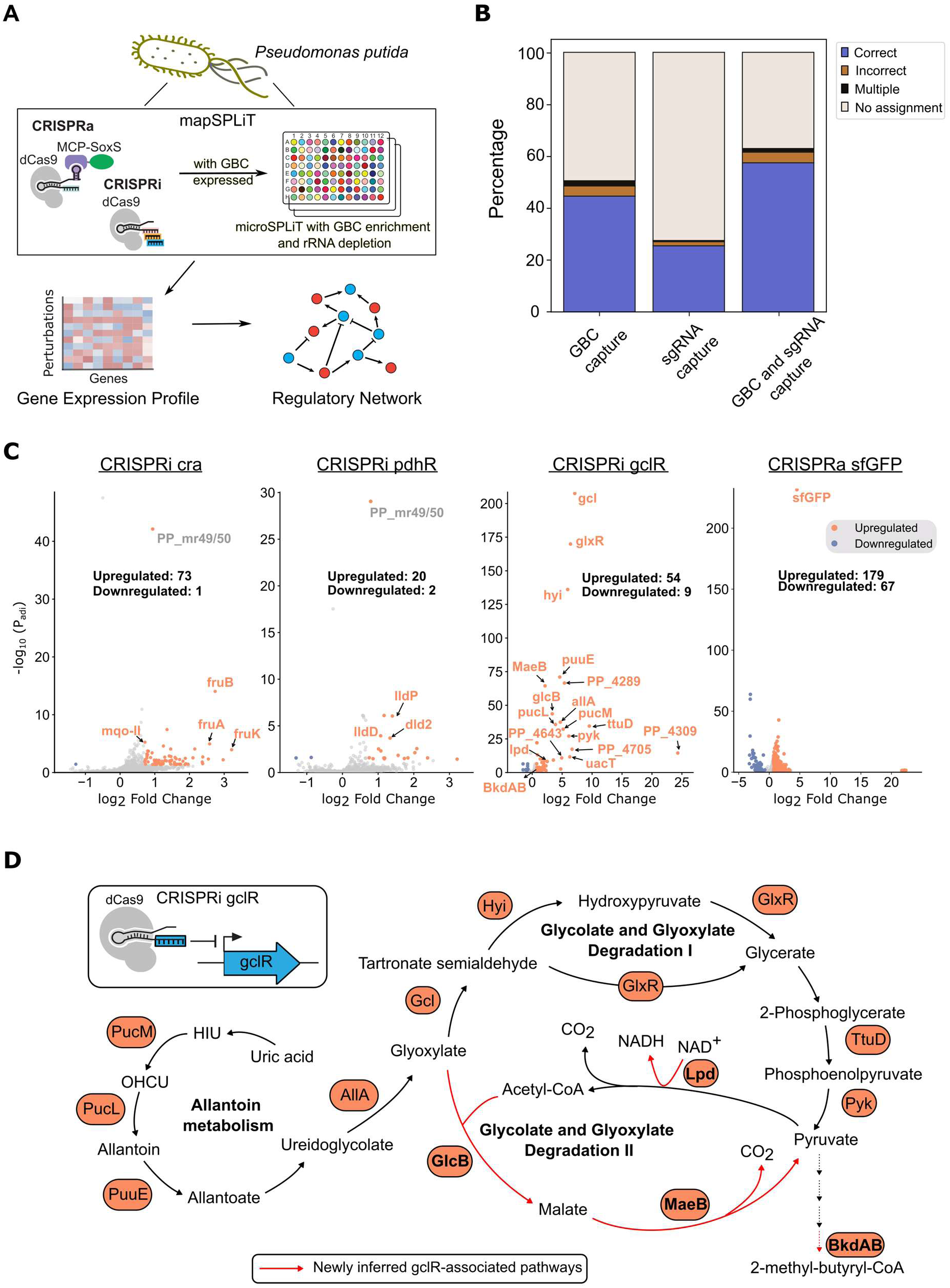
mapSPLiT reveals transcriptional networks in *P. putida.* **a**, The established mapSPLiT workflow is ported to *P. putida* unchanged. **b,** Percentage of cells assigned to perturbations grouped by: correctly assigned, incorrectly assigned, expressing multiple GBCs/sgRNAs, or unassigned. Cells are assigned using only GBC capture, only sgRNA capture, or both. **c,** All captured genes (gray points) are plotted with their expression (x-axis) and p-values from the Wilcoxon rank sum test (y-axis). DEGs are plotted in orange. Genes previously annotated or putatively assigned to the regulon of the perturbation target are labeled in orange. For cra and pdhR perturbations, genes with p-values < 10⁻²⁵ are labeled in gray. **d,** Metabolic pathways associated with the GclR regulon. DEGs (orange) from the CRISPRi perturbation targeting *gclR* are mapped onto the allantoin utilization and glycolate and glyoxylate degradation I/II pathways. Newly inferred GclR-associated pathways are shown in red, and their corresponding genes are bolded.

Next, we determined the regulatory responses for each CRISPRi perturbation by analyzing their DEGs. (**table S8, Methods**). We examined 7,509 cells assigned to perturbations with a median of 82 transcripts and 66 genes per cell (**fig. S17**). We confirmed that mapSPLiT can detect CRISPRa responses in *P. putida*, as the CRISPRa perturbation of *sfGFP* upregulated expression 4.5-fold (**Fig. 6C**). Next, we analyzed the transcription factor perturbations. We recapitulated many regulatory connections established in previous literature^49–53,55^ when perturbing *cra*, *pdhR*, and *gclR* with CRISPRi (**Fig. 6C**). We discovered that *cra* regulates a malate:quinone oxidoreductase in the TCA cycle (*mqo-II*), consistent with its predicted binding site^49^. This finding supports the idea that, similar to its *E. coli* counterpart, *P. putida cra* functions as a global regulator extending beyond fructose metabolism. CRISPRi of *gclR* led to upregulation of allantoin pathway genes, glyoxylate degradation II genes, a gene linking the two glyoxylate degradation pathways, and downstream genes shared by both pathways (**Fig. 5, C and D**). These differential expression responses align with predicted regulatory connections^11,56^ or prior proteomics data^57^. Because these pathways enable assimilation of nitrogen-rich purines and C₂/C₃ intermediates, their coordinated regulation suggests that *gclR* plays a broader role in carbon and nitrogen metabolism than currently annotated, with implications for feedstock utilization. Collectively, these results showcase that mapSPLiT can be seamlessly adapted to *P. putida*, enabling measurement of both CRISPRa and CRISPRi responses to discover regulatory relationships.

## Discussion

Single-cell CRISPR screens have transformed functional genomics in eukaryotes. In bacteria, sparse transcript capture has prevented scalable coupling of pooled perturbations with single-cell sequencing. We developed mapSPLiT, a pooled CRISPRa/i screening platform that integrates guide barcode PCR enrichment and sequencing-based rRNA depletion into the microSPLiT framework. These advances allow mapSPLiT to profile single-cell transcriptomes and link them to perturbations in the same cells across hundreds of pooled perturbations. For the first time, bacterial CRISPR perturbation responses can be resolved at genome-wide scale with single-cell RNA sequencing. mapSPLiT can be scaled along two axes: number of single cell transcriptomes, which increases exponentially with microSPLiT combinatorial barcoding rounds^4,58^, and gene targets, by building larger sgRNA libraries. Together, these features create new capacity to interrogate gene activities, detect genetic interactions, and reconstruct regulatory circuits in bacteria.

Applied to *E. coli*, we scaled mapSPLiT to over a hundred transcription factor perturbations in a single experiment. This application expanded known regulatory networks, assigned functions to putative regulators, and revealed new direct connections. Transcriptome-wide profiling of 76,000 *E. coli* cells showed that many perturbations converged on shared global states while others by themselves produced multiple distinct phenotypes. In *P. putida*, differential gene expression analysis uncovered an expanded role for the transcriptional regulator *gclR* in coordinating glyoxylate and allantoin metabolism. These findings suggest that tuning *gclR* could act as a genetic “switch” for unconventional substrate assimilation, enhancing *P. putida* as a chassis to convert lignin-derived aromatics, waste plastics, and nitrogen-rich residues into high-value chemicals, fuels, and bioplastics^59^. Together, these results demonstrate that single-cell CRISPR screens expose unknown biology and deliver systems-level insights into complex regulatory architectures.

mapSPLiT uncovered unexpected genetic interactions when multiple genes were targeted simultaneously in the same cell. Multi-gene perturbations produced non-additive outcomes, including synergistic and buffering effects not predictable from individual contributions. The *aroF*/*tyrR* perturbation pair exemplified this pattern, generating widespread non-additive responses across the transcriptome. In contrast, *tyrR* and *trpR* directed expression of their co-regulated targets through additive outcomes governed by a dominant regulator. These findings show that bacterial transcriptional networks exhibit a spectrum of genetic interactions resulting in complex regulatory logic, similar to recent findings in other bacterial systems^60,61^.

Advancing microbial engineering requires generalizable technologies for systematic exploration of genome-wide functions^62^. By linking perturbation genotypes to single-cell transcriptome phenotypes, mapSPLiT generates data that can inform models of bacterial metabolism and regulation^63,64^. The scale of these datasets enables integration with AI/ML approaches to uncover hidden patterns and predictive features^65^. Together, these capabilities establish a foundation for more predictive microbial biosystems design, with applications spanning the bioeconomy and biomedicine^66,67^. We anticipate that mapSPLiT will open new avenues of inquiry across disciplines, paralleling the transformative impact of analogous eukaryotic platforms.

## Methods

### Experimental methods

#### Bacterial strain construction and manipulation

Plasmid constructs expressing perturbation-specific sgRNAs and GBCs were individually cloned using a common sgRNA vector, expressing a chloramphenicol resistance gene, dCas9, and MCP-SoxS. The vector was digested with BcuI and PacI (NEB) and gene fragments containing the unique 12 bp barcode and sgRNA spacer sequences for each perturbation (IDT) were inserted with In-Fusion Snap Assembly (Takara Bio). Resulting plasmids were transformed into chemically-competent DH5α *E. coli*. Each member of the library was then isolated and sequence-verified separately then transformed into *E. coli*. MG1655 to create our final strains. Where indicated, strains were pooled and grown together. In all experiments, the strains were grown overnight in EZ-RDM (0.2% glucose and chloramphenicol) while shaking at 37°C in a 250 mL flask. Following overnight incubation, cultures were diluted 1:250 in 12 mL of media in 15 mL tubes and grown as before until OD_600_=0.4. In the high-throughput pooled experiment, we used a vector with inducible dCas9. dCas9 expression was induced with 5 nM aTc for the CRISPRa pool and 200 nM aTc for the CRISPRi pool during the second growth step.

#### Microbial single-cell RNA sequencing using split-pool ligation transcriptomics

After growing strains as described in “Bacterial strain construction and manipulation,” we performed the standard microSPLiT protocol as described by Gaisser et al.^58^ and Kuchina et al.^8^. Where noted, modifications were made to the protocol to include “Guide barcode construct enrichment” and “Sequencing-guided rRNA depletion.” Final cDNA libraries were either sequenced with Illumina Nextseq 500 (150 cycles) or sent to Novogene for pre-made library sequencing on Illumina platforms.

#### Guide barcode construct enrichment

Before the on-bead cDNA amplification (step 105 in Gaisser et al.^58^), 9.68 µL of a GBC-specific primer (oJRB099: 5’-GCGGATGTAGGATGGTCTCCAGACAC-3’ at 100 µM) was added at a ratio of 5:1:1 (oJRB099:BC_0108:BC_062) alongside standard primers. BC_62 served as the reverse primer for the whole transcriptome and the GBC enrichment.

#### Sequencing-guided rRNA depletion

We designed 253 sgRNA spacer sequences to target the 5S, 16S, and 23S rRNA sequences in *E. coli* and *P. putida*, weighted proportionally to the regions in the rRNA that appeared more frequently during NGS after microSPLiT. These spacer sequences were ordered in an oligo pool (IDT) with the necessary 5’ and 3’ overhangs for *in vitro* transcription using the EnGen® sgRNA Synthesis Kit (NEB) according to manufacturer’s protocol. 100 pmoles of the transcribed, column-purified sgRNA molecules were pre-incubated with 10 pmoles spCas9 (NEB) in a 30 μL with NEB r3.1 Buffer at room temperature for 10 minutes. We then added 30 ng of the mapSPLiT cDNA library (from after the final double-sided size selection, step 171 in Gaisser et al.^58^) to the pre-incubated reaction for a final volume of 45 μL, then incubated at 37°C for 2 hours. The reaction containing the depleted cDNA library was cleaned with KAPA Pure Beads (Roche) at 1X, qPCR-amplified using P5 and P7 adapter primers (Illumina), then cleaned again with KAPA Pure Beads at 0.7X before sequencing as described in “Microbial single-cell RNA sequencing using split-pool ligation transcriptomics.” For comparisons between rRNA-depleted and rRNA-undepleted sequenced libraries (**Fig. 2**) both libraries were downsampled to the same number of raw reads.

#### D-lactate colorimetric assay

Single colonies of strains were seeded in 3 mL EZ-RDM (0.2% glucose and chloramphenicol) and grown overnight then subcultured as described in “Bacterial strain construction and manipulation.” The D-lactate assay (MilliporeSigma) was performed on cultures at varying growth states (OD_600_ 0.4, 0.7, and 1.5) and measured by a microplate reader (Biotek Synergy HTX). The assay was performed and the data were analyzed according to the manufacturer’s protocol. The output of the assay for each sample was normalized to the strain’s growth state (OD_600_). To determine statistical significance relative to an off-target control, a two-sided Welch’s t-test was used.

#### Plate reader experiments

Single colonies of strains were seeded into 96-deep-well plates with 400 µL EZ-RDM (0.2% glucose and chloramphenicol) while shaking at 37°C for 22 hours. 150 µL of overnight cultures were transferred into a 96-well clear flat-bottom plate and the fluorescence was measured by a microplate reader (Biotek Synergy HTX) set to measure sfGFP (excitation: 485 nm, emission: 528 nm, gain: 35). To determine statistical significance for a sample relative to an off-target control, a two-sided Welch’s t-test was used.

#### Quantitative RT-PCR

Single colonies of strains were seeded in 3 mL EZ-RDM (0.2% glucose and chloramphenicol) and grown overnight then subcultured as described in “Bacterial strain construction and manipulation”. These cultures were spun down and RNA was extracted using an RNeasy Mini Kit (QIAGEN). Reverse transcription was performed with random hex primers (IDT) and Maxima H Minus Reverse Transcriptase (ThermoFisher). A quantitative PCR was run with KAPA HiFi HotStart (Roche) and EvaGreen (Biotium) on a CFX Opus Dx Real-Time PCR System (Bio-Rad), which also determined C_t_ values. Relative expression was calculated using 2^-^_ΔΔCt._

### Computational methods

#### Choosing CRISPR perturbation targets

In the high-throughput pooled experiment (**Fig. 3**), we chose to perturb 27 “well-characterized” and 24 “partially characterized” transcription factors, totalling 51 transcription factor targets. The “well-characterized” transcription factors were chosen to encompass a broad range of regulon sizes, from local pathway regulators to broad global regulators, and biological processes. Some “partially characterized” transcription factors were chosen because they had putative binding sites from prior work^68,69^ while others were randomly selected. We also included CRISPRa perturbations of shikimate kinase I (*aroK*) to further test CRISPRa on genes in aromatic amino acid biosynthesis (**fig. S10C**), bringing the total number of on-target gene targets to 52.

#### Guide barcode construct design

The GBC was designed as a 316 nt non-coding RNA transcript such that with cell barcodes and Illumina adapters attached would correspond to the size range selected for sequencing (∼450 nt). Each GBC includes a unique 12 nt barcode, an enrichment primer (JRB099) binding site upstream of the barcode, an Illumina adapter binding site upstream of the barcode, and a 3’ polyadenylation sequence. The bulk of the 316 nt transcript consisted of a portion of the *Lachnospiraceae* bacterium dCas12a gene, chosen to be efficiently transcribed but not translated. The length was chosen to ensure that the entire GBC was appropriate for sequencing. The unique 12 nt barcodes were designed using the dna-barcodes Python tool (v1.4.12)^70^ with exactly 50% GC content (**table S2**).

#### Single-guide RNA design

To design custom on-target CRISPRi sgRNAs for genomic perturbations we first selected all possible targets on the non-template strand within the first 250 bp of the start of the open reading frame. For on-target CRISPRa sgRNAs, which have stringent rulesets for bacteria^10,66^, we first selected all possible targets on both the template and non-template strand within 2 bp of the optimum distances from the transcription start site (70, 80, 90 bp). We further screened these sgRNAs for performance by using the kinetic folding algorithm described in Fontana et al.^71^. We chose all sgRNAs that had a bind barrier < 10 kcal/mol and a new net energy < -25 kcal/mol. We tiled two sgRNAs per targeted gene if target site requirements and folding parameters allowed.

For off-target, negative control sgRNAs, we designed their spacer sequences by randomly generating 12 spacer sequences with 50% GC content using a custom Python script. Sequences with binding complementarity (less than 5 mismatches) to the *E. coli* genome predicted by the RGEN Cas-OFFinder tool^72^ were removed, and the resulting sequences were filtered through the kinetic folding algorithm described in Fontana et al.^71^ with the same thresholds as the on-target sgRNAs.

#### Alignment and generation of cell-gene matrices from single-cell data

The STARsolo software (v2.7.10a)^73^ was used to align cDNA reads to a custom reference genome (ASM584v2 for *E. coli* or ASM756v2 for *P. putida* with sequences dCas9, MCP-SoxS, all sgRNAs, and all GBC transcripts for the experiment), deduplicate reads, and create a count matrix for downstream processing. All further processing of the single-cell data was performed using Scanpy (v1.9.3)^74^. To determine our filtering thresholds, we followed standard scRNA-seq practices by filtering out cells with less than a threshold number of transcripts as determined by the inflection point of the cell barcode rank plot. All rRNA and tRNA genes were then removed. We then referenced a second cell barcode rank plot to perform a second filtering. We retained cells that had greater than a threshold number of genes (3 genes per cell) and retained genes that had greater than a threshold number of transcripts (3 transcripts per gene in *E. coli* or 5 transcripts per gene for *P. putida*). Counts were then normalized, log-transformed, and scaled as described in the Scanpy documentation^74^.

#### Perturbation assignment

To assign cells in our dataset to a specific perturbation, we used either only the expression of GBCs, only the expression of sgRNAs, or both, where indicated. We assigned each cell to a perturbation using the most highly-captured transcript (GBC or sgRNA) that aligned to that perturbation. If multiple transcripts had equal capture rates and were the most highly-captured transcripts, we labelled the cell as having “Mulitple” assignments. Cells in which no relevant transcripts (GBC or sgRNAs) were captured were not assigned to any perturbation. All multiple assigned and unassigned cells were filtered out prior to further analyses.

#### Differential gene expression analysis

Following “Alignment and generation of cell-gene matrices from single-cell data” and “Perturbation assignment,” we performed a Wilcoxon rank sum test with a tie correction. When performing the Wilcoxon, we compared the expression profile of each individual on-target perturbation to all of its corresponding CRISPRa or CRISPRi off-target controls, treated as a single sample for CRISPRa or CRISPRa. We set a stringent threshold to reduce the likelihood of false positives and defined DEGs for each perturbation as any gene that had p_adj_<0.05 and abs(log_2_-fold change) > 0.7 from the Wilcoxon rank sum test. As an increase in the proportion of cells can indicate enhanced gene expression^75^, we further removed DEGs that were expressed in less than 5% of cells. The order of the perturbations in the heatmap (**Fig. 4C**) was determined by using a dendrogram. All of these analyses were done in Scanpy (v1.9.3)^74^.

#### Gene regulatory network analysis

For each perturbation targeting a transcription factor, we classified its DEGs as either a result of a direct regulatory network connection or an indirect regulatory network connection. We labeled DEGs as “direct regulatory network connection” if the gene belonged to the regulon of the targeted transcription factor according to the EcoCyc database^29^ (or by using previously-published transcription factor binding experiments^68,69^ for our putative transcription factor targets). All other DEGs were labeled as “indirect regulatory network connection.” We could further annotate any “direct regulatory network connection” as an activation or repression depending on the direction of change in the expression of the DEG.

#### Genetic interaction analysis

For sets of perturbations, we calculated genetic interaction scores for every DEG in at least one of the perturbations. For each DEG, we calculated the mean log_2_-fold change (relative to an off-target control) in the combined sgRNA perturbation (“measured”) and compared it to the additive sum of the log_2_-fold changes in all single sgRNA perturbations (“additive”). We subtracted the additive from the measured log_2_-fold changes to calculate the genetic interaction score. For three-guide programs, we compared both the pairwise combination of sgRNAs separately and all three together.

To determine which genetic interaction scores were statistically significant, we used the 95% confidence interval of bootstrapped scores. Confidence intervals for the mean log_2_-fold changes and the gene interaction scores were determined by a bootstrap analysis. Using a custom Python script, bootstrapped mean log_2_-fold changes and gene interaction scores for each gene and for each single and combined perturbation were calculated as above by resampling sets of cells with replacement over 1000 iterations. We considered genetic interaction scores to be significantly non-additive if the 95% confidence interval of the bootstrapped genetic interaction scores did not pass through 0.

To determine whether the difference in mean transcripts per cell at the *tyrR* and *trpR* co-regulated genes *aroL* and *mtr* was statistically significant (**Fig. 3H, fig. S8B**), we determined the empirical p-value^76^ of the distribution of the bootstrapped transcripts per cell. The null hypothesis was defined as the mean transcripts of the on-target perturbation being the same or less than that of the off-target perturbation. The empirical p-value was computed as the fraction of all pairwise comparisons in which a value drawn from the on-target bootstrap distribution was less than or equal to a value drawn from the off-target bootstrap distribution.

#### Gene set enrichment analysis

Gene set enrichment analyses were performed with the GOATOOLS python package^77^. For each perturbation, each DEG was classified as either upregulated and downregulated. The enriched gene ontology (GO) terms were evaluated for the up- and downregulated genes separately. The GO terms were filtered (p<0.05 using a two-sided Fisher’s exact test) then ranked by p-value. The top 15 of these GO terms were then selected by significance and iteratively filtered. For each parent-child GO term pair in this list, the parent GO terms were removed, then another GO term was added from the initial ranked list. This process was repeated until no parent-child GO term pairs remained.

#### Single-cell clustering to identify perturbation-induced cell states

We first filtered the scRNA-seq data to only include DEGs that were significant in at least one perturbation using a more stringent threshold than in “Differential gene expression analysis” (p_adj_<0.05, abs(log_2_-fold change) > 1, expressed in > 5% of cells with that perturbation). Then, we performed dimensionality reduction on our dataset using principal component analysis and UMAP^34^. We plotted the neighborhood graph with 20 principal components (PCs), chosen from the plateau of the plot of the variance ratio for the top 50 PCs, and chose 10 neighborhood points. The dimensionality was further reduced to 2 dimensions using UMAP. We then manually annotated the resulting clusters using the marker genes identified by the leiden algorithm (with a resolution of 0.5). All of these analyses were done in Scanpy (v1.9.3)^74^ and according to scRNA-seq best practices as described in Luecken et al.^78^.

We then identified perturbations enriched in each leiden cluster by first calculating the percentage of the perturbation’s cells that belonged to that cluster. We set a threshold of either 10% of cells from the perturbation or one standard deviation greater than the percentage of cells from all off-target perturbations in that cluster, whichever was greater. Any perturbation that passed this threshold was considered enriched for a specific cluster (**table S7**). Enriched perturbations were plotted in UMAP space by taking the mean X and Y coordinates of all cells with that perturbation in each cluster.

## Author contributions

JRB, QT, YH, DS, and KDG designed and performed experiments and analyzed data. JRB, JMC, and AK prepared the original draft. JRB, QT, YH, GS, JGZ, JMC, and AK contributed to writing and revision. GS, JGZ, JMC, and AK conceived and supervised the study and secured funding. All authors approved the final manuscript.

## Acknowledgments

We thank members of the Zalatan, Seelig, Carothers and Kuchina groups for advice. The authors used an AI assistant (Microsoft Copilot) to provide feedback on grammar and refine sentence structure; all content was reviewed, edited, and approved by the authors, who take full responsibility for the final publication. The information, data, or work presented herein was funded in part by the U.S. Department of Energy under Award Number DE-SC0023091. This report was prepared as an account of work sponsored by an agency of the United States Government. Neither the United States Government nor any agency thereof, nor any of their employees, makes any warranty, express or implied, or assumes any legal liability or responsibility for the accuracy, completeness, or usefulness of any information, apparatus, product, or process disclosed, or represents that its use would not infringe privately owned rights. Reference herein to any specific commercial product, process, or service by trade name, trademark, manufacturer, or otherwise does not necessarily constitute or imply its endorsement, recommendation, or favoring by the United States Government or any agency thereof. The views and opinions of the authors expressed herein do not necessarily state or reflect those of the United States Government or any agency thereof.

## Data availability

All relevant sequencing files will be deposited to the Sequence Read Archive (SRA) at the time of publication. Processed data will be submitted to GEO at the time of publication. Scripts used for data analysis and visualization will be made available on GitHub. Annotated plasmid sequence files will be made available at the time of publication.

