## Supplementary information for "Pooled single-cell CRISPRa/i screens for functional genomics in bacteria at scale"

### Supplementary Text

Supplementary note 1: Efficacy of GBC enrichment and direct sgRNA capture in *E. coli*

To investigate the impact of enrichment primer concentration on perturbation assignment, we compared two concentrations of enrichment primers relative to the whole transcriptome primers (2X, 5X). Without any PCR enrichment of the GBC, we recovered 4869 cells and were able to assign 5% of them to the correct perturbation by the detected GBC sequence. With a GBC enrichment primer present at 2X concentration relative to the transcriptome primers, we assigned 16% of cells to the correct perturbation, 2.9-fold higher than the no enrichment set. To determine if assignment could be further improved with a higher primer ratio, we increased the GBC primer to 5X concentration and observed that correct assignment increased to 21%, 3.8-fold higher than no enrichment (**Fig. 1C**).

To investigate whether PCR-based enrichment of the GBC led to barcode swapping events that produced spurious GBC-cell barcode pairings, we examined the expression of all GBCs across assigned cells. It has been previously demonstrated that increasing the number of PCR cycles during the library preparation process leads to more uncoupling between sgRNAs and their barcodes[1]. Therefore, as higher concentrations of enrichment primer could be equivalent to additional cycles of PCR, we reasoned that there could be uncoupling between the GBCs and their associated cell barcodes. For both GBC primer enrichment ratios, there was no evidence for substantial barcode swapping as most cells expressed a single GBC (0.4% multiple assigned) and the incorrect assignment percentage did not increase across enrichment conditions (<2%) (**Fig. 1C**). We also investigated whether competition between the GBC enrichment and cDNA amplification PCR reactions impacted the recovery of cellular transcriptomes. Total transcripts per assigned cell (not including GBCs) decreased on average by 29% for the 5X condition and by 16% for the 2X condition relative to no GBC enrichment (**Fig. 1C**). Thus, with increasing levels of GBC primer, there is a tradeoff between improved GBC assignment and total transcript recovery, and any further increases in the GBC enrichment primer ratio are likely to further decrease transcriptome recovery. We also observed that although there was a decrease in transcript recovery with enrichment, the cells that were successfully assigned tended to have higher levels of detectable transcripts (**fig. S1A**). To improve mapSPLiT cost-efficiency and throughput, we proceeded with a 5X GBC enrichment primer ratio for all subsequent experiments, reasoning that the 3.8-fold increase in assignment outweighed the decrease in transcriptome recovery.

We evaluated the efficacy of GBC assignment relative to using only the sgRNA and found that only 8-10% of cells could be correctly assigned if the GBC is ignored (**fig. S1B**). To improve assignment with sgRNAs, we evaluated two possible approaches. First, we considered PCR enrichment of the sgRNA itself. In practice, however, this approach would be impractical because each sgRNA contains a unique spacer sequence at the 5’ end, which would require a distinct enrichment primer for every individual sgRNA. Second, we considered combining GBC and sgRNA assignment together. For GBC with no enrichment or 2X primer enrichment, including sgRNA resulted in improved assignment relative to either GBC or sgRNA alone (**fig. S1B**), with the 2X GBC and sgRNA condition producing relatively high assignment (>20%). However, there was an ~67% increase in incorrectly assigned cells. This increase is potentially problematic because we aim for high confidence in assignment calls to perform pooled screens. For GBC with 5X primer enrichment, including sgRNA did not increase assignment, presumably because the GBC assigned all cells that had sufficiently high transcript counts. Across all conditions tested, the 5X enrichment GBC condition produced both high correct assignment (>20%) and comparably low incorrect assignment, and we proceeded with the 5X enrichment GBC condition for all future experiments.

### Supplementary note 2: Impact of CRISPRa-mediated upregulation of *aroL* and *poxB*

In the CRISPRa *poxB* strain, we did not detect any significant gene expression changes for the *poxB* target (**fig. S2D**) or any downstream genes (**Fig. 1E**). We confirmed with RT-qPCR in the same conditions (see Methods) that our CRISPRa *poxB* strain did effectively increase the expression of *poxB* by 4.5-fold (**fig. S3A**). Despite this upregulation, *poxB* expression still remained very low at $2^{20}$-fold less than 16S rRNA in RT-qPCR (**fig. S3B**) and $7x10^{-4}$ transcripts per cell in mapSPLiT (**fig. S3C**), so we reasoned that *poxB* expression was below the limit of detection in our experiments.

For the *aroL* CRISPRa perturbation, we did not directly observe activation of the target gene (**fig. S2D**). Unlike the transcription factors targeted for CRISPRi above, *aroL* expression levels are high enough that sparse coverage is unlikely to explain the lack of *aroL* upregulation (**fig. S4A**). However, it is possible that the *aroL* gene was upregulated initially but then a regulatory feedback response from *tyrR* or *trpR* reduced expression of the gene. Nonetheless, we observed transcriptional changes in several other metabolic genes that could be affected by changes in *aroL* levels and corresponding effects on aromatic amino acid bioproduction (**fig. S4, B and C**). Specifically, we observed downregulation of glycolytic genes such as *fbaA*, *pykA*, *pfkA,* and *ppc* (**fig. S4C**). Similar effects were previously observed in an *E. coli* strain engineered to overproduce phenylalanine[2]. These gene expression changes are a plausible regulatory response to upregulation of the aromatic amino acid bioproduction pathway. Downregulation of glycolytic genes should decrease phosphoenolpyruvate (PEP) availability, which would drive metabolic flux away from amino acid bioproduction. Additionally, we found that tyrosine and phenylalanine tRNA synthetase genes were upregulated (*pheT*, *pheS*). These synthetases attach free tyrosine and phenylalanine to their corresponding tRNAs. It is plausible that as tyrosine and phenylalanine are products of the aromatic amino acid biosynthesis pathway that *aroL* was upregulated at some point during the growth process. We also observed upregulation of translation machinery, consistent with increased protein demand (**fig. S4D**). Thus, the CRISPRa system is likely activating *aroL* as expected.

### Supplementary note 3: Evaluating the sequencing-guided rRNA depletion method

Our rRNA depletion method resulted in significant depletion of rRNA reads and an enrichment of mRNA reads in the sequenced library. Downsampling to the same number of raw reads, we observed a decrease in the proportion of rRNA UMIs from 94.6% to 20.1%, and an increase in the proportion of mRNA UMIs from 4.6% to 60.2% (**Fig. 2B**). After filtering our single-cell data (see Methods), the sequencing-guided rRNA depletion method resulted in a final dataset with a median of 88 mRNA genes per cell and 115 mRNA UMIs per cell in the rRNA-depleted sample compared to 49 mRNA genes per cell and 54 mRNA UMIs per cell in the undepleted sample (**Fig. 2C**). We wanted to further verify that rRNA depletion did not lead to differential capture of any non-rRNA transcripts and resulted in the same mRNA expression profiles. Comparing the capture of all transcripts between the depleted and undepleted samples showed high correlation for both mRNA and tRNA (Pearson r = 0.983 and 0.981, respectively) (**Fig. 2D**). Additionally, the capture of the most highly-expressed rRNA genes (23S rRNAs) was reduced by a mean of 12.6-fold, bringing them on par with the expression of highly-expressed non-rRNA genes (**Fig. 2D**). This confirms that our sequencing-guided cleavage sgRNA library presents a cost-effective solution to effectively reduce rRNA reads in the cDNA sublibrary without affecting non-rRNA reads. We used this rRNA depletion method for all subsequent mapSPLiT experiments.

<u>Supplementary note 4: Pathway-level interactions for the CRISPRa of *aroF* and CRISPRi of *tyrR* perturbation pair</u>

To interrogate pathway-level genetic interactions from the *aroF/tyrR* perturbation, we performed hierarchical clustering for all mRNA transcripts that met our significant gene threshold in the *aroF*/*tyrR* perturbation pair (**Fig. 3D**). We observed buffering of glycolysis genes (*pykA*, *gapA*, *pfkA*, *pgk*) characteristic of a feedback response to reduce metabolic flux into aromatic amino acid biosynthesis. This behavior is consistent with downregulation of the same genes observed in a phenylalanine overproducing strain[2]. We also observed buffering of fermentation (*adhE*, *pflB*) and anaerobic metabolism or low oxygen conditions genes (*frdA*, *frdB*, *cydB*, *focA*, *aspA*) which could also limit carbon intake. Finally, we observed synergistic activation of electron transport genes (*cyoE*, *cyoB*, *fdoG*) and an associated oxidative stress response (*sodA*). Taken together, the genetic interactions from *aroF/tyrR* perturbations reveal connections to both glycolysis and electron transport, suggesting potential regulatory connections between aromatic amino acid metabolism and multiple steps in metabolism. This data is consistent with previous work demonstrating that changes in metabolic flux alter the activity of the transcription factor, *cra*[3], which controls genes in both of these pathways[4].

<u>Supplementary note 5: Regulatory interactions at the *aroL* and *mtr* promoters</u>

At the *mtr* promoter, repression of *tyrR* alone led to a modest, statistically insignificant decrease in expression compared to an off-target control ($p>0.05$). Conversely, we observed that repressing *trpR* alone resulted in a 4-fold increase in *mtr* expression compared to an off-target control ($p<0.001$), suggesting that *trpR* acts as a repressor (**fig. S8B**). However, repressing both *tyrR* and *trpR* increased *mtr* expression only 2.4-fold compared to the off-target control ($p<0.001$), suggesting that *tyrR* functions as an activator when *trpR* repression is absent. Our initial genetic interaction analysis (**Fig. 3, G and F**) did not identify a significant non-additive interaction between *tyrR* and *trpR* within the 95% confidence interval that we applied for statistical significance. Instead, this interaction appears within a 91% confidence interval. Our initial analysis was stringent to avoid false positives at the risk of false negatives. Overall, these results suggest that there could be a genetic interaction between *tyrR* repression and *trpR* activation at the *mtr* promoter, and that *trpR* repression dominates the regulation of *mtr* expression.

At the other co-regulated promoter, *aroL*, we observed that the CRISPRi perturbation targeting only *tyrR* increased *aroL* expression compared to an off-target control ($p<0.001$, 1.7-fold change) whereas CRISPRi targeting *trpR* alone had no effect ($p>0.05$, 0.88 fold-change) (**Fig. 3H**). Simultaneously perturbing both *tyrR* and *trpR* has the same effect as perturbing *tyrR* alone, suggesting that *tyrR* alone regulates *aroL* expression. We verified this conclusion by performing a fluorescence reporter assay with the same sets of perturbations and observed the same pattern of *aroL* responses (**Fig. 3H**). Overall, despite prior expectations that *tyrR* and *trpR* both contribute to *aroL* expression, our data suggest that *tyrR* is the sole regulator of *aroL*. At both co-regulated promoters, our results largely agree with previous work[5–7]. However, there are some inconsistencies that could be explained by differences in growth condition, assay sensitivity, or variations in transcription factor concentrations from knockdown compared to knockouts. These differences could be systematically explored in future mapSPLiT experiments across many conditions in parallel.

Supplementary note 6: Newly discovered regulatory connections across the library of CRISPR perturbations

All regulators introduced below are "partially-characterized" other than the *cpxR* transcription factor. We target some regulators with pre-existing transcription factor binding data allowing for partial inference of their function while others had no experimental data at all. In our study, we observed the first expression data for 6 regulators without previous binding data (*ytfA*, *ytfH*, *phnF*, *lgoR*, *sgcR*, *sfsB*). Regardless of prior data, we found new connections for 17 regulators. We describe the responses to *yheO* perturbation in the main text, while the rest are highlighted below.

*ytfA*

Although annotated as a pseudogene, CRISPRi perturbation of *ytfA* increased expression of the known cell division gene *dapE*[8], and the known biofilm formation regulator *bssS*[9]. Previous data showing that *ytfA* mRNA is upregulated in a biofilm-deficient *tqsA* knockout[10], further corroborates a functional role for the *ytfA* gene product.

*ytfH*

We observed that knockdown of *ytfH*, previously inferred as a stress-responsive transcription factor[11], results in a large increase in *qorB* gene expression (albeit in ~4% of the cells). As *qorB* was exclusively differentially expressed in the *ytfH* perturbation, the magnitude of gene expression was large ($\log_2$-fold change of 4.6), and expression was approaching the cell threshold, we included CRISPRi of *ytfH* in our analysis (**Fig. 4, A, B, D**). *qorB*, which has been implicated in reducing quinone toxicity and antibiotic resistance[12], is located immediately upstream and in the opposite orientation of *ytfH*. *ytfH* could regulate *qorB* by divergent transcription (e.g., *ilvY* regulation of *ilvC*[13]), connecting a putative role of *ytfH* in stress adaptation to quinone detoxification (Supplementary figure 10e).

*yjhI*

We observed upregulation of *yjhH* and *yjhG* in less than 5% of the cells for one of the two CRISPRi perturbations we performed (**fig. S10D**). We suspect that *yjhI* regulates *yjhH* and *yjhG* but dCas9 CRISPRi polar effects resulted in lower than expected expression of the downstream genes in the regulatory network.

*plaR*

*plaR*, is a repressor of a single operon whose leader gene is *yiaK*. As this operon has been annotated as requiring derepression of *plaR* and activation by *crp*, we suspected that we would not observe significant activation without perturbing both transcription factors. However, we did observe expression of *yiaK* in less than 5% of the perturbations' cells (**fig. S10D**).

*decR*

*decR* is an activator of L-cysteine import (*cyuP*, *cyuA*) that we targeted with CRISPRi. Although we did not detect upregulation of the lowly expressed regulatory network genes, we did observe a strong upregulation of a sulfite reductase, *cysJ*, which is a key enzyme involved in L-cysteine biosynthesis from L-serine (**fig. S10D**). A genome-wide SELEX study confirmed that the *cyuPA* operon is the only regulatory target of *decR*[14]. Therefore, we uncovered a cellular response to diminished cysteine concentrations from the loss of function of *decR*.

*cpxR*

The two component envelope stress response cascade is composed of the sensor histidine kinase *cpxA* and the transcription factor *cpxR*. We validated previously known regulatory network connections when perturbing *cpxR* with CRISPRi but also captured expression of *yagU* and *raiA* which are induced at stationary phase or due to various stress responses. These genes have been previously determined to be in the *cpxR* i-modulon[15], but the expression of *yagU* and *raiA* has not been demonstrated by direct knockout or knockdown.

*yciT*

The *yciT* regulator controls genes involved in osmolarity and has prior transcription factor binding and RNA-seq validation using ChIP-exo[16]. Our analysis shows a similar gene expression profile to ChIP-exo. However, we also observed expression of the *dmsABC* operon which was revealed in ChIP-seq but not the RNA-seq validation.

*yddM*

Upon knockdown of the *yddM* gene, we observed a broad downregulation of motility and chemotaxis genes consistent. Notably, we detected downregulation of the serine sensing chemotaxis protein, *tsr*, which was previously identified as a target of *yddM* using transcription factor binding data[17]. We surmised that loss of a motility sensor attenuated nutrient sensing leading to reduced expression of flagellar genes.

*yfiE*

We found that perturbing *yfiE* resulted in upregulation of the glycerol degradation V pathway enzymes: *gldA*, *dhaL* and *dhaK*. Additionally, transcription factor binding[16] experiments identified *yfiE* binding sites within the *dhaKLM* operon suggesting that *yfiE* directly regulates the transcription of *dhaL* and *dhaK*. These enzymes are part of a complex that converts dihydroxyacetone to glycerone phosphate. The conversion of glycerol to dihydroxyacetone is mediated by *gldA*, which may be indirectly regulated by *yfiE*. More broadly, there were over 300 hundreds DEGs impacted by perturbing *yfiE* with CRISPRi. We categorized these upregulated genes by GO terms and found biological processes such as cell motility, anaerobic respiration, regulation of cell shape, cadmium ion transport, and glycerol-3-phosphate catabolic process enriched (**Fig. 4D, fig. S12**). This data suggests that *yfiE* has a global regulatory role beyond glycerol metabolism.

*phnF*

The *phnCDEFGHIJKLMNOP* operon is responsible for organophosphonate utilization as part of the carbon-phosphorus lyase pathway[18]. We find that repression of *phnF*, a putative transcriptional regulator based on structural studies[19], led to differential expression of the small regulatory RNA-expressing gene *glmY* and *maeA* which codes for malate dehydrogenase. Because CRISPRa/i modulates only the transcription of a gene, the repression of other genes in the polycistronic *phn* transcript may contribute to the effect we see for CRISPRi of *phnF*.

*lgoR*

*lgoR* has very little prior characterization; *lgoR* expression is regulated by and required for the metabolism of L-galactonate as the sole carbon source in *E. coli* growth[40]. In our screen, CRISPRi of *lgoR* resulted in the differential expression of *fliA*, *fliC*, *rimK*, *bioB*, *yjiA*, *mgtA*, and *rnr*. Although none of these genes pertain to L-galactonate, this is likely because regulators *uxuR* and *exuR* repress *lgoR* unless alpha-D-glucuronate is present[20]. Therefore, a focused multi-guide mapSPLiT experiment could uncover the interactions between *lgoR* and the upstream transcriptional regulators on the downstream L-galactonate genes (*lgoT*, *lgoD*).

*sgcR, sfsB, ybcM, yfeC, ybaQ*

For each of these regulators, we observed limited gene expression information or broad transcriptional responses such as the expression of motility genes. Additional mapSPLiT experiments with multiple sgRNA targets for each of these regulators could elucidate further regulatory connections.

### Supplementary note 7: *ldhR* regulation of pyruvate flux under aerobic and anaerobic conditions

We confirmed that *ldhR* regulates *ldhA* using single-cell transcriptomics, a fluorescent reporter assay, and a colorimetric metabolite assay. Previous work has demonstrated that *ldhA* is primarily active under anaerobic conditions[21], but the measurable D-lactate produced by the off-target control indicates that some D-lactate is synthesized aerobically. The second target of *ldhR* is *lipA,* which encodes a lipoyl synthase that attaches lipoate to pyruvate and 2-oxoglutarate dehydrogenases[22]. By regulating *lipA*, *ldhR* could balance pyruvate flux between D-lactate production and central metabolism under aerobic conditions. Under anaerobic conditions, when pyruvate dehydrogenase is inactive, flux would instead be directed to D-lactate production. This evidence is supported by the observation that the off-target control is still producing measurable D-lactate under aerobic conditions.

Supplementary note 8: Regulatory convergence in the Histidine biosynthesis cluster

Knockdown of *pdhR* with CRISPRi produced polar effects on downstream operon genes encoding pyruvate dehydrogenase (*aceE*, *aceF*, *lpd*) which could cause a bottleneck in pyruvate consumption. We observed the activation of a general stress response (*dnaK*, *clpB*, *groL*, *grpE*, *dnaJ*, *lon*, *htpG*, *hslU*, *hslV*, *hslO*, *groS*, *recA*), as well as the with upregulation of genes involved in NADH production or utilization (*gapA*, *aldA*, *hisD*, *nuoM*) and downregulation of NADPH-consuming sulfur assimilation genes (*cysD*, *cysN*, *cysJ*, *cysK*, *tcyJ*, *tcyP*, *cysP*), a pattern that could reflect a regulatory shift to compensate for altered NADH and NADPH. Therefore, downregulation of *pdhR* may cause upregulation of histidine biosynthesis as a consequence of disrupting central metabolism.

Supplementary note 9: Efficacy of GBC enrichment and direct sgRNA capture in *P. putida*

To perform GBC enrichment, we employed the same unmodified GBC sequence and added the enrichment primer at a 5X concentration which in *E. coli* resulted in the highest correct assignment of cells to perturbations. For all CRISPRa and CRISPRi perturbations, we evaluated cell assignment using both the GBC and the sgRNA. Using solely the GBC, we found that correct assignment was 44.6% (**Fig. 6B**), 2.1-fold higher than *E. coli*, indicating that the GBC was more highly expressed, the GBC was more stable, or the enrichment PCR was more efficient in *P. putida*. Furthermore, we found that using both the GBC and sgRNA sequence yielded 57.4% correct assignment (**Fig. 6B**), 1.3-fold higher than GBC alone and 2.3-fold higher than the sgRNA alone. These results indicate that the sgRNA enhances assignment efficiency and that the GBC and sgRNA capture distinct populations of cells. Unlike in *E. coli*, including sgRNA for assignment in *P. putida* did not substantially increase incorrect assignment compared to GBC assignments alone.

Supplementary note 10: Evaluating the sequencing-based Cas9 rRNA depletion protocol in *P. putida*

We found that rRNA comprises ~95% of reads in *P. putida* mapSPLiT data (**fig. S16A**). Motivated by reducing the associated costs of sequencing rRNA, we implemented the sequencing-guided rRNA depletion method that we developed for *E. coli*. Using the same raw number of reads, the proportion of rRNA transcripts decreased from 95.2% to 73.0% and the proportion of mRNA transcripts increased from 2.2% to 10.4% (**fig. S16A**). Of note, the increase in the proportion of mRNA was lower in *P. putida* (4.7-fold) (**fig. S16A**) than in *E. coli* (13-fold) (Figure 2b), consistent with previous work showing that the efficacy of Cas9-based rRNA depletion varies across bacteria[23]. Nevertheless, our strategy improved the resolution of our *P. putida* dataset (**fig. S16B**) and did not affect the sequencing of non-rRNA genes (**fig. S16C**). Accordingly, we used the rRNA-depleted dataset for the differential gene expression analysis.

Overall, our depletion strategy proved transferable to *P. putida* and further optimization could likely enhance its efficacy further.

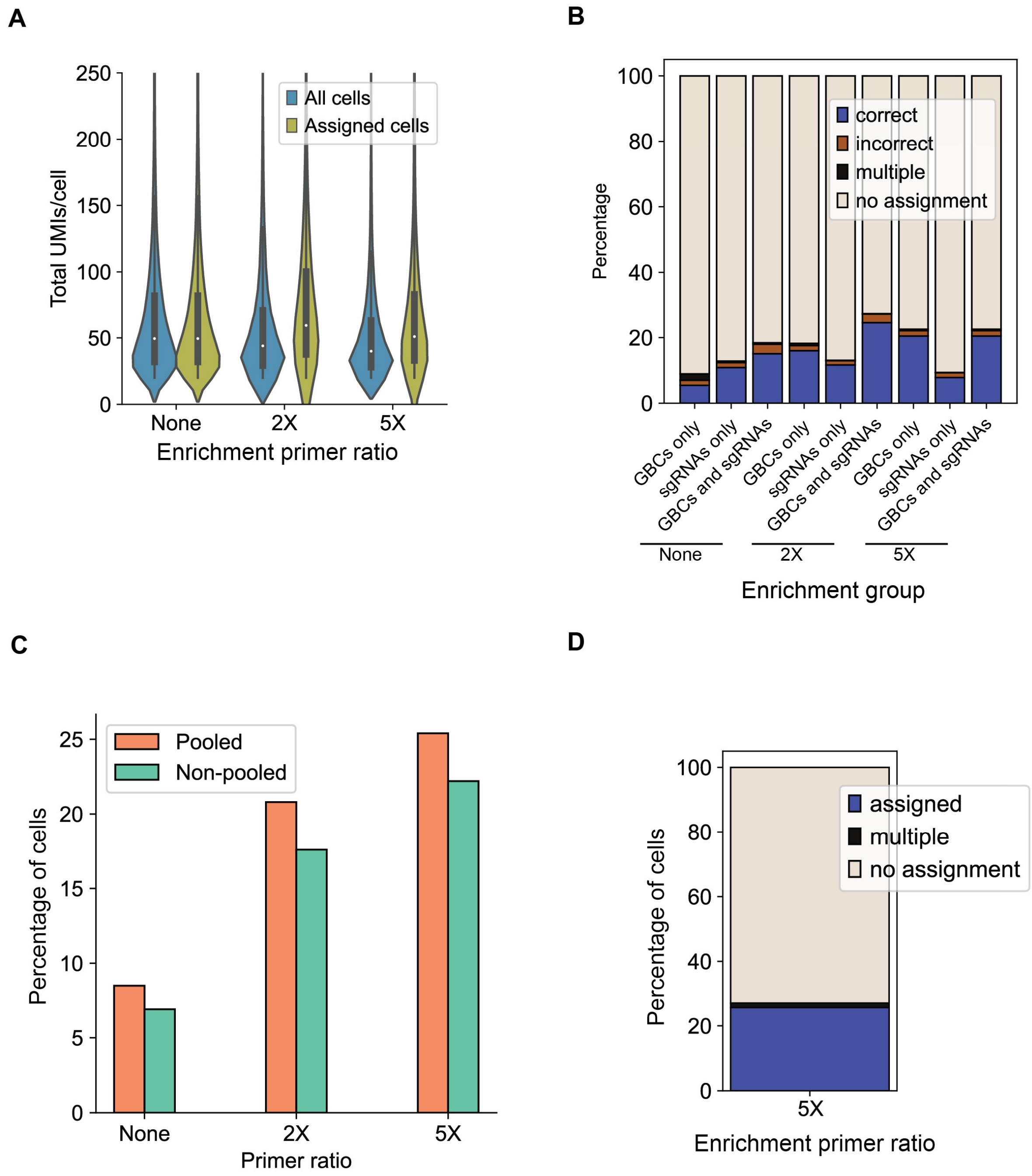


**Fig. S1. Optimizing guide barcode construct enrichment. (A)** Distribution of total transcripts per cell using the GBC (different concentrations of enrichment primer relative to reverse primer) for cells before CRISPR perturbation assignment and after. **(B)** Percentage of assigned cells using the GBC (different concentrations of enrichment primer relative to reverse primer), the sgRNA, or both by category (correct, incorrect, multiple, or no assignment). **(C)** Percentage of assigned cells for the pooled and arrayed cells using the GBC as in **(A)**. **(D)** The percentage of assigned cells by category as in **(B)** utilized for transcriptional network analysis.

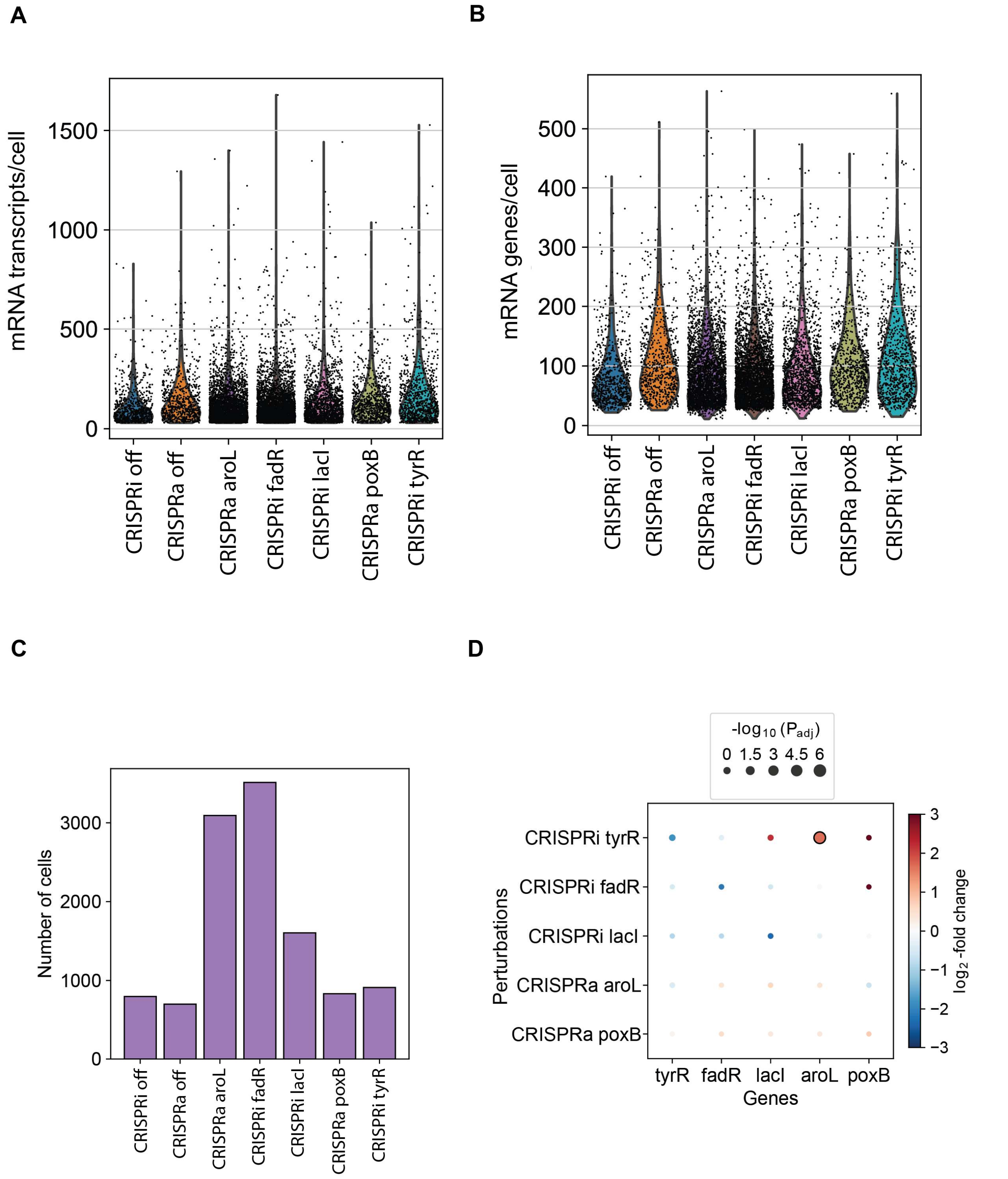


**Fig. S2. Single-guide single-cell library sequencing metrics. (A)** Distribution of genes per cell with each dot representing a single cell. **(B)** Distribution of genes per cell with each dot representing a single cell. **(C)** Cell counts across CRISPR perturbations. **(D)** Expression of the target genes for each CRISPRa and CRISPRi perturbation. The dot sizes represent the adjusted p-values from the Wilcoxon rank sum test relative to an off-target control. Outlines indicate statistical significance.

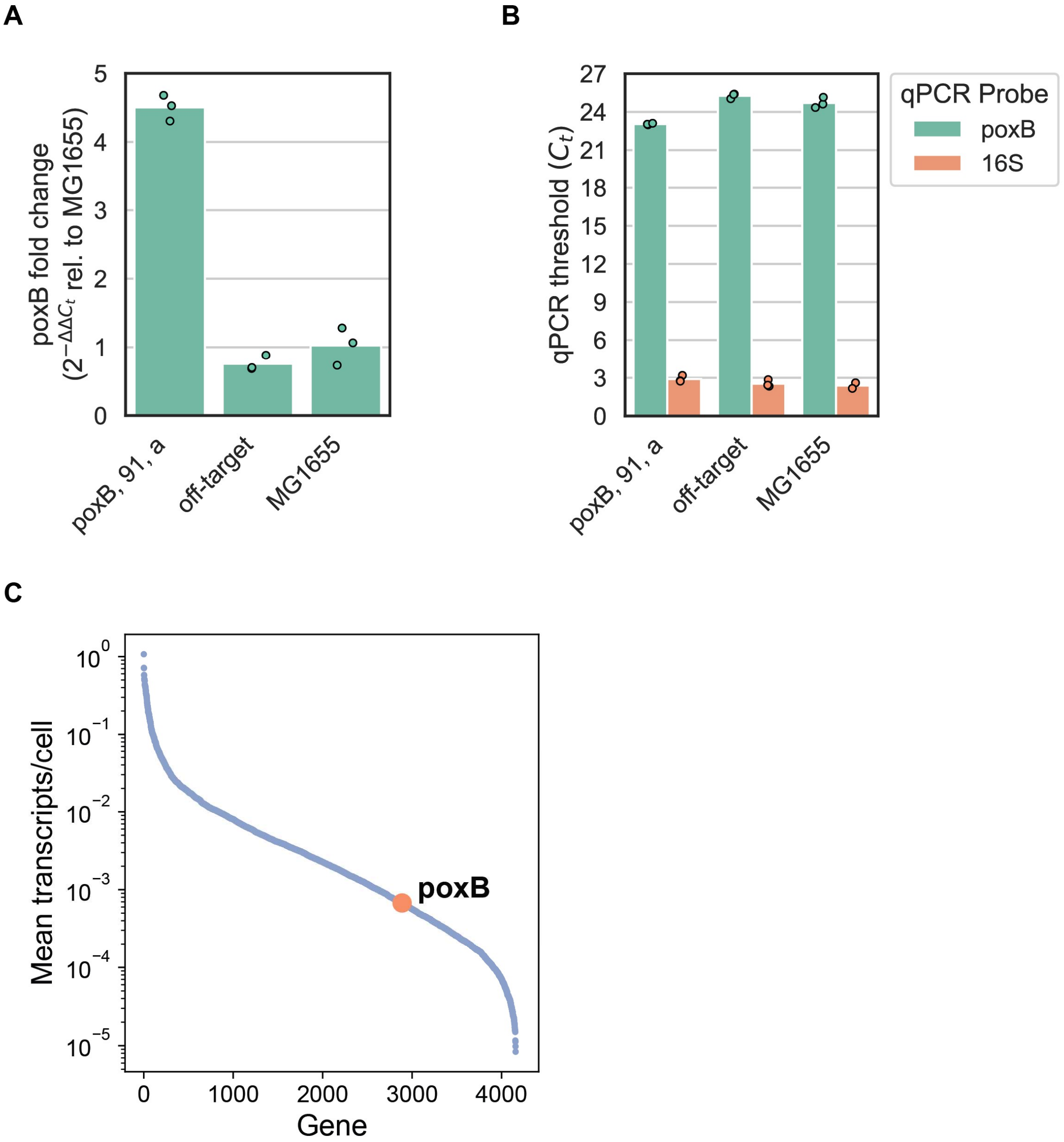


**Fig. S3. Evaluating the *poxB* CRISPRa response in bulk and single-cell RNA sequencing.** **(A)** Relative expression of poxB ($2^{-\Delta\Delta C_t}$ relative to 16S expression and in MG1655) of a poxB CRISPRa, a CRISPRa off-target, and a wild-type MG1655 strain. **(B)** Ct values of a *poxB* CRISPRa, a CRISPRa off-target, and a wild-type MG1655 strain for *poxB* and 16S qPCR probes (as described in Fontana et al.). Ct values were determined by CFX Opus Dx Real-Time PCR System (see Methods). **(C)** Mean transcripts per cell for all *E. coli* genes (blue) with the dot (orange) corresponding to the *poxB* gene.

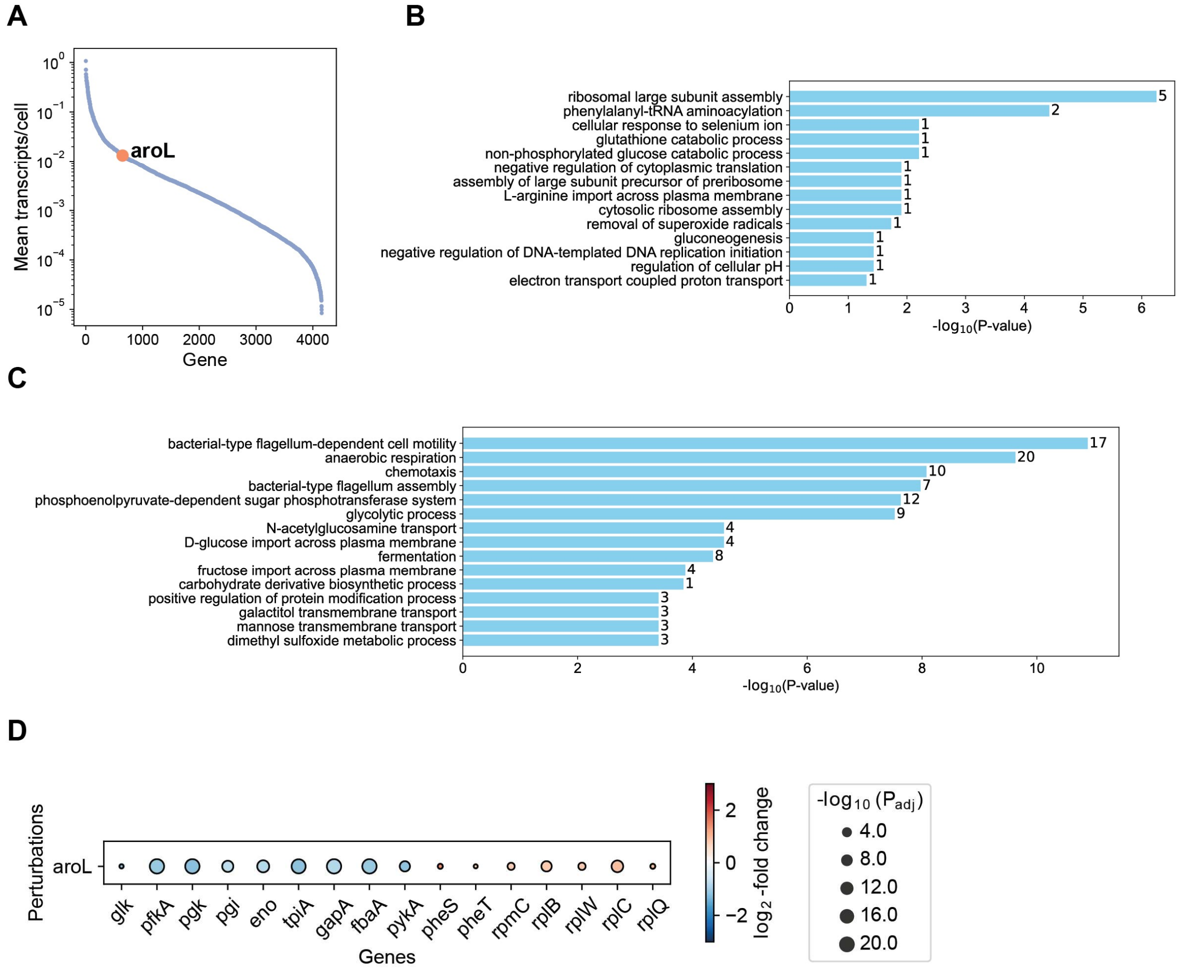


**Fig. S4. CRISPRa of *aroL* transcriptional responses in the single guide pilot experiment.** **(A)** Mean transcripts per cell for all *E. coli* genes (blue) with the dot (orange) corresponding to the *aroL* gene. **(B)** GO-term enrichment of select biological processes calculated with the upregulated DEGs for the *aroL* CRISPRa perturbation. The p-values were calculated using a two-sided Fisher's exact test. **(C)** As in (a) using the downregulated DEGs. **(D)** Expression of glycolytic, tRNA, and translational genes identified from the GO-term enrichment analysis. The size of dots represent the adjusted p-values from the Wilcoxon rank sum test relative to an off-target control. Outlines indicate statistical significance.

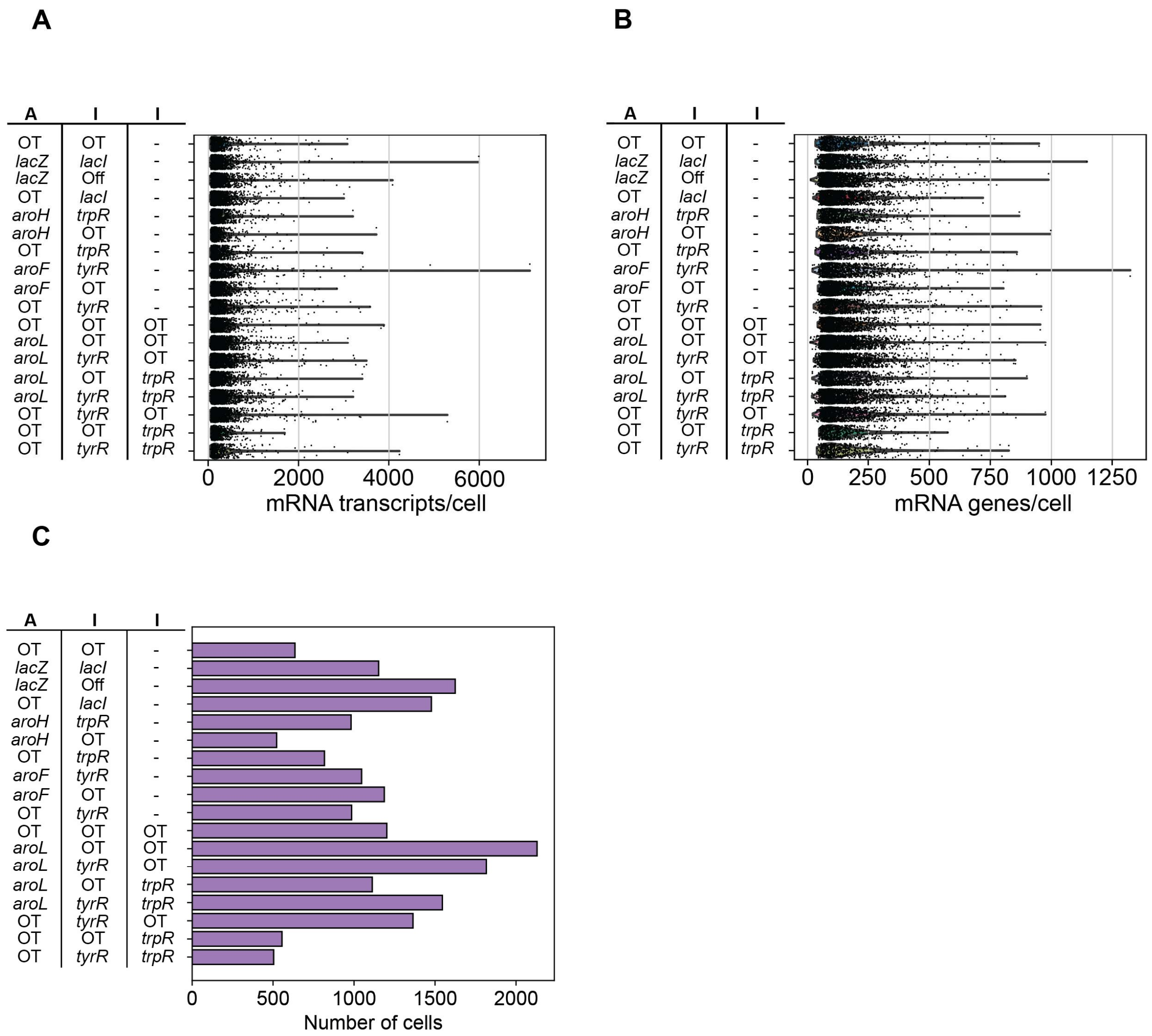


**Fig. S5. Multi-guide single-cell library sequencing metrics. (A)** Distribution of transcripts per cell with each dot representing a single cell. Each row is a separate strain with A indicating CRISPRa and I indicating CRISPRi. The dash ("-") indicates a 2-guide and not a 3-guide strain. "OT" represents an off-target sgRNA. (**B)** Distribution of genes per cell with each dot representing a single cell. Each row is as in **(A)**. **(C)** Cell counts across CRISPR perturbations. Each row is as in **(A)**.

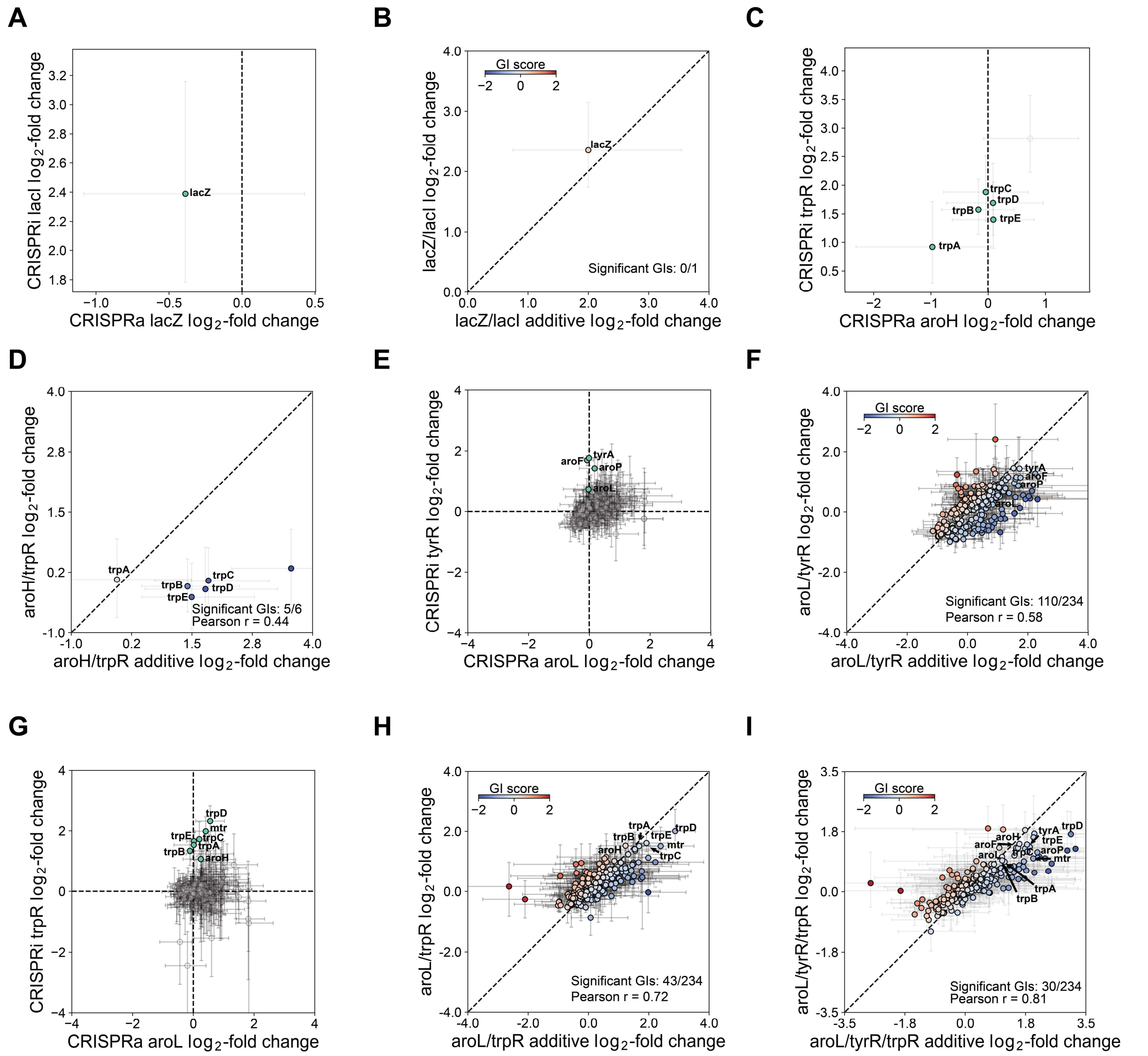


**Fig. S6. Computing genetic interactions (A, C, E, G, and I)** Single sgRNA $\log_2$-fold changes of DEGs for CRISPRa of *lacZ* and CRISPRi of *lacI* **(A)** CRISPRa of *aroH* and CRISPRi of *trpR* **(C)** CRISPRa of *aroL* and CRISPRi of *tyrR* **(E)** CRISPRa of *aroL* and CRISPRi of *trpR* **(G)** perturbations relative to an off-target control. Error bars are bootstrapped 95% confidence intervals (CIs). Green dots and text indicate CRISPRa target gene or regulon of targeted transcription factor. **(B, D, F, H, and I)** Genetic interaction map for CRISPRa of *lacZ* and CRISPRi of *lacZ* **(B)** CRISPRa of *aroH* and CRISPRi of *trpR* **(D)** CRISPRa of *aroL* and CRISPRi of *tyrR* **(F)** CRISPRa of *aroL* and CRISPRi of *trpR* **(H)** CRISPRa of *aroL*, CRISPRi of *tyrR*, and CRISPRi of *trpR* **(I)** depicting the measured (x-axis) and predicted additive (y-axis) $\log_2$-fold change for each DEG relative to an off-target control. Error bars are as in **(A, C, E, G, I)**. Colorbar shows the genetic interaction (GI) score as the difference between the measured and predicted $\log_2$-fold changes. Text indicates CRISPRa target gene or regulon of targeted transcription factor.

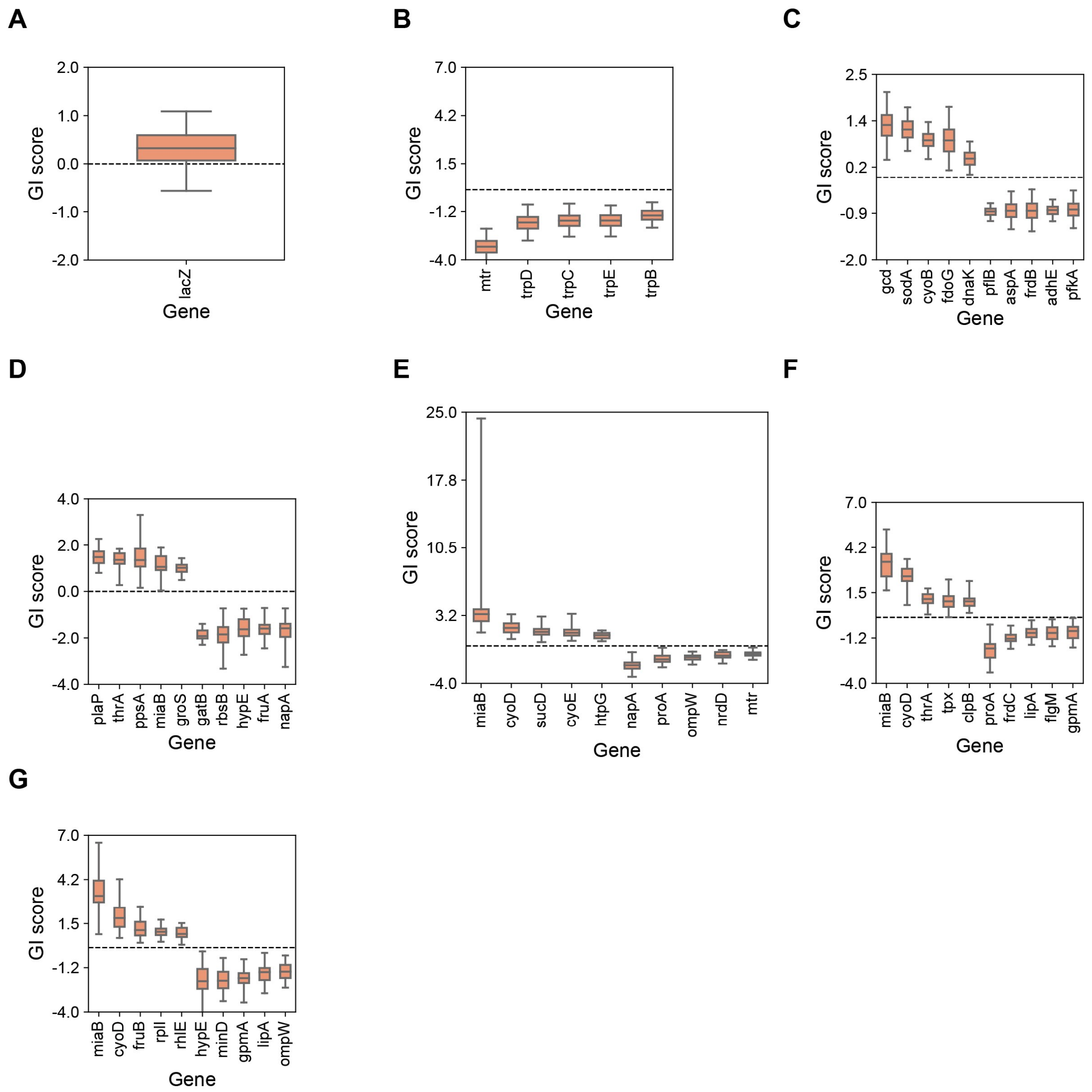


**Fig. S7. Genetic interactions scores (A, B, C, D, E, F, and G)** Up to 10 significant GI scores (top 5 synergistic and top 5 buffering by absolute score) for CRISPRa of *lacZ* and CRISPRi of *lacI* **(A)** CRISPRa of *aroH* and CRISPRi of *trpR* **(B)** CRISPRa of *aroF* and CRISPRi of *tyrR* **(C)** CRISPRa of *aroL* and CRISPRi of *tyrR* **(D)** CRISPRa of *aroL* and CRISPRi of *trpR* **(E)** CRISPRi of *tyrR* and CRISPRi of *trpR* **(F)** CRISPRa of *aroL*, CRISPRi of *tyrR*, and CRISPRi of *trpR* **(G).**

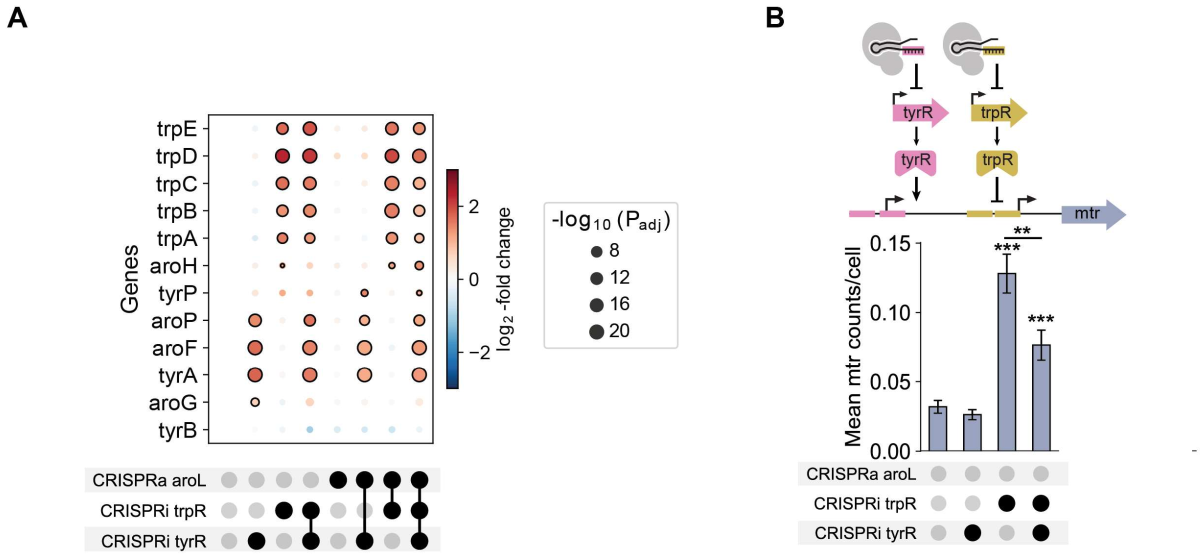


**Fig. S8. *tyrR* and *trpR* regulon. (A)** Expression of genes solely regulated by *tyrR* or *trpR*. The size of dots represents the adjusted p-values from the Wilcoxon rank sum test relative to an off-target control. Outlines indicate statistical significance. **(B) h,** Top, schematic of CRISPRi sgRNAs targeting *tyrR* and *trpR* at the *mtr* promoter. Bottom, mean *mtr* transcript counts per cell with bootstrapped standard deviations (SDs). Empirical p-values (see Methods), *p<0.05, **p<0.01, ***p<0.001. Black circles indicate contributing perturbations.

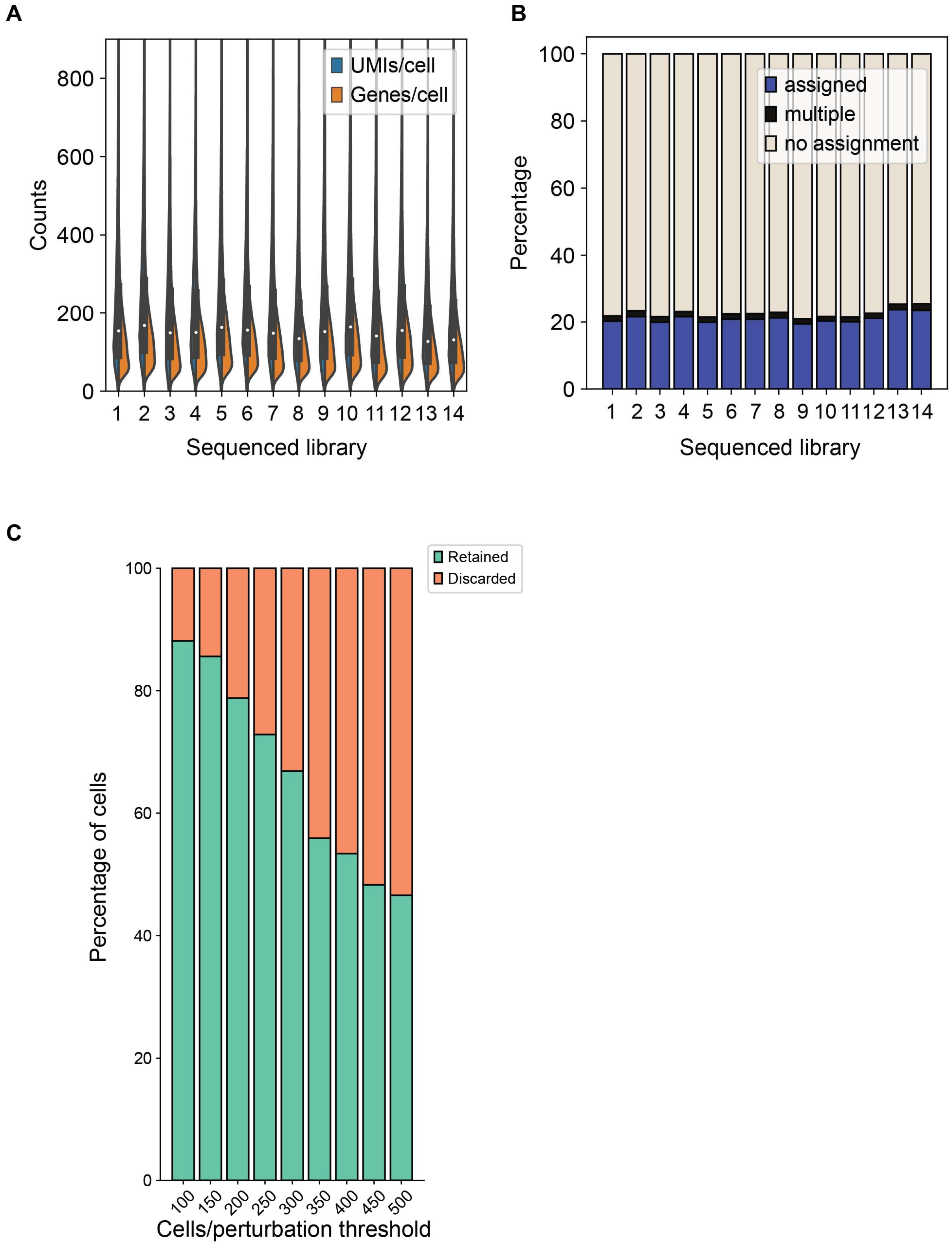


**Fig. S9. High-throughput single-cell library sequencing metrics. (A)** Split violin plots depicting transcripts/cell (left) and genes/cell (right) for 14 sequenced libraries prior to perturbation assignment. **(B)** Percentage of cells assigned for all 14 sequenced libraries prior to perturbation assignment. **(C)** Percentage of cells retained when requiring perturbations to have a minimum number of cells.

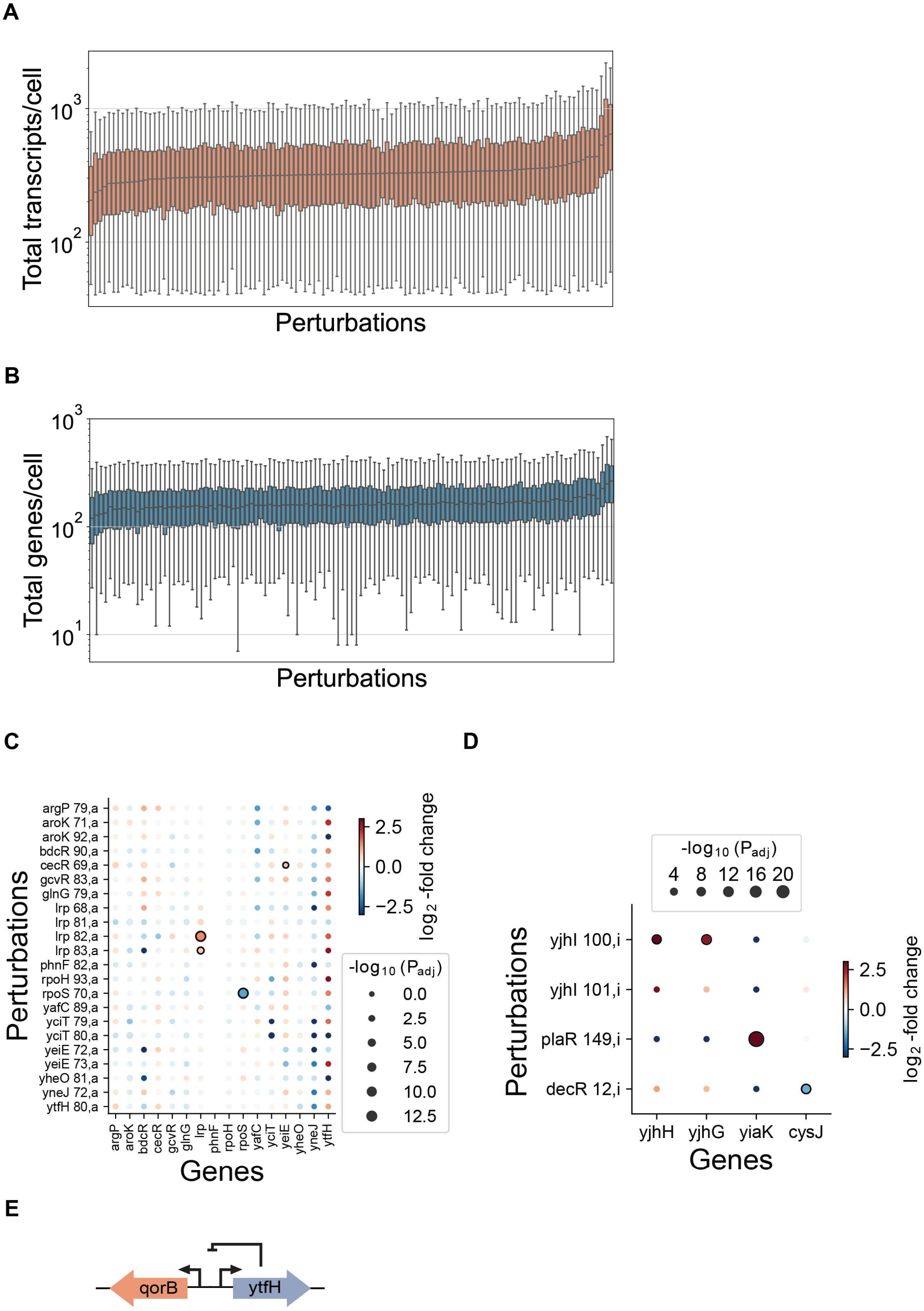
A
Total transcripts/cell
$10^3$
$10^2$
Perturbations
B
Total genes/cell
$10^3$
$10^2$
$10^1$
Perturbations
C
argP 79,a
aroK 71,a
aroK 92,a
bdcR 90,a
cecR 69,a
gcvR 83,a
glnG 79,a
lrp 68,a
lrp 81,a
lrp 82,a
lrp 83,a
phnF 82,a
rpoH 93,a
rpoS 70,a
yafC 89,a
yciT 79,a
yciT 80,a
yeiE 72,a
yeiE 73,a
yheO 81,a
yneJ 72,a
ytfH 80,a
Perturbations
argP
aroK
bdcR
cecR
gcvR
glnG
lrp
phnF
rpoH
rpoS
yafC
yciT
yeiE
yheO
yneJ
ytfH
Genes
2.5
0.0
−2.5
$log_2$-fold change
$-log_{10}$ ($P_{adj}$)
0.0
2.5
5.0
7.5
10.0
12.5
D
$-log_{10}$ ($P_{adj}$)
4 8 12 16 20
yjhI 100,i
yjhI 101,i
plaR 149,i
decR 12,i
Perturbations
yjhH
yjhG
yiaK
cysJ
Genes
2.5
0.0
−2.5
$log_2$-fold change
E
qorB
ytfH

**Fig. S10. Characterizing the library of perturbations. (A)** Median and IQRs of total transcripts per cell ordered by median from low to high for each assigned perturbation. **(B)** Median and IQRs of total genes per cell ordered by median from low to high for each assigned perturbation. **(C)** The differential expression of the CRISPRa perturbations on the on-target genes. The size and outlines of the dots are as in **(C)**. **(D)** Expression of DEGs that are expressed in less than 5% of a perturbation's cells. The dot sizes represent the adjusted p-values from the Wilcoxon rank sum test relative to an off-target control. Outlines indicate statistical significance. **(E)** Schematic showing the proposed model of *ytfH* regulation on itself and *qorB* based on transcriptomic data.

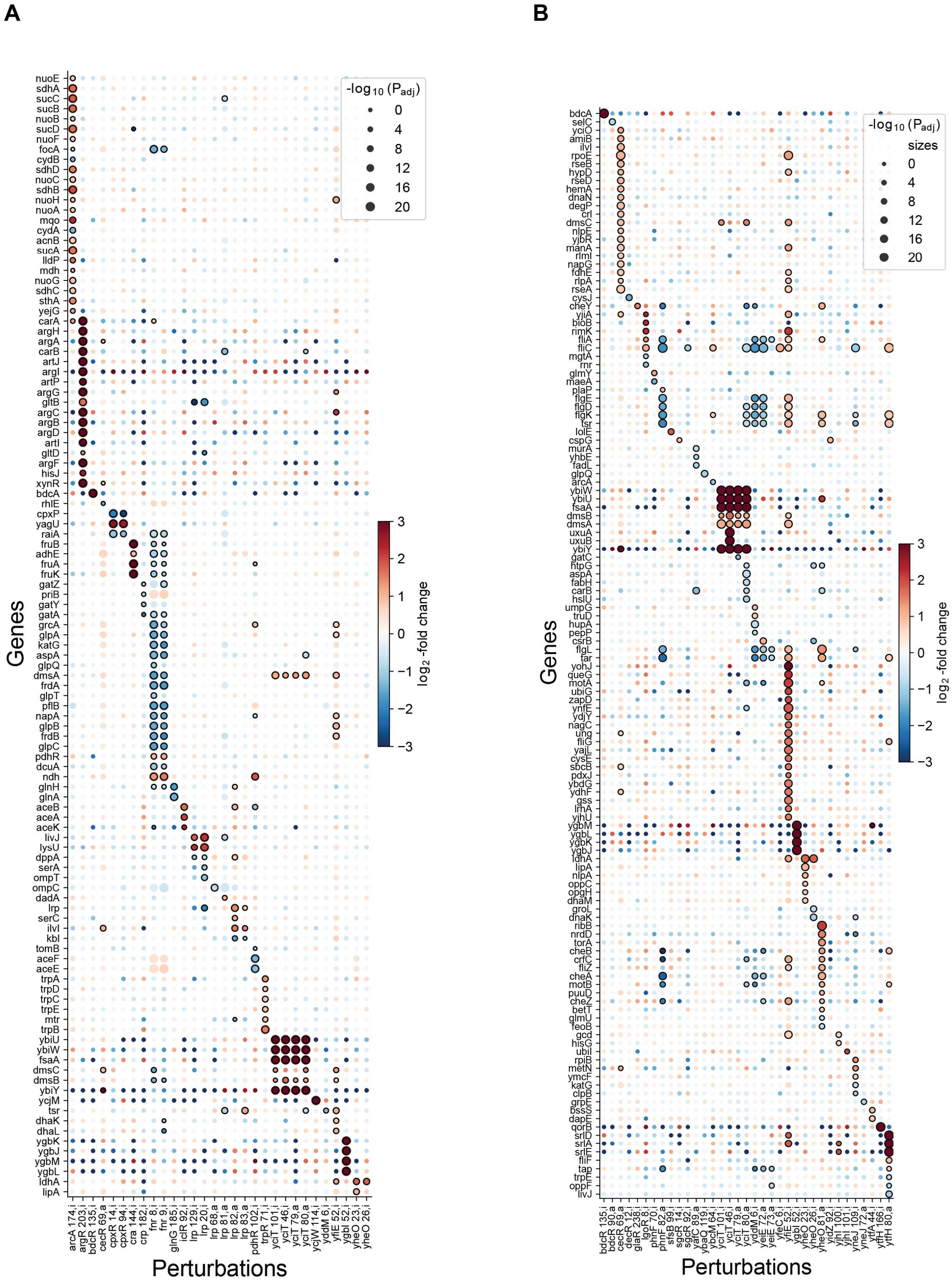
A
B
Genes
Perturbations
-log$_{10}$ (P$_{adj}$)
sizes
0
4
8
12
16
20
log$_2$ -fold change
3
2
1
0
−1
−2
−3

**Fig. S11. Measuring transcriptional responses for a library of perturbations. (A)** Expression of DEGs for perturbations with prior transcription factor binding data. All included genes are binding partners of at least one transcription factor. The dot sizes represent the adjusted p-values from the Wilcoxon rank sum test relative to an off-target control. Outlines indicate statistical significance. **(B)** As in **(A)** for the top 20 DEGs ranked by $\log_2$-fold change for the “partially characterized” transcription factor perturbations.

**A**

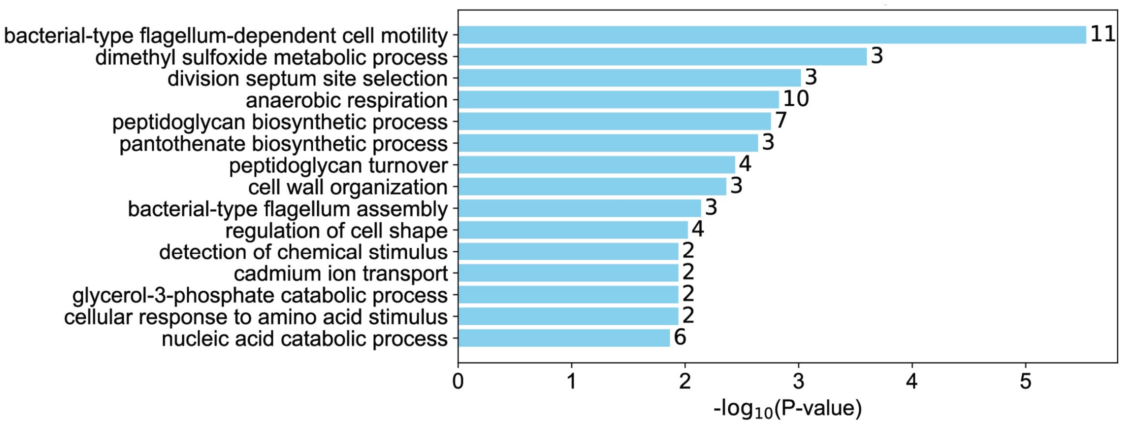


**Fig. S12. Impact of CRISPRi of *yfiE* on biological processes. (A)** GO-term enrichment of select biological processes calculated with the upregulated DEGs for the *yfiE* CRISPRi perturbation (this perturbation only resulted in upregulated DEGs). The p-values were calculated using a two-sided Fisher's exact test.

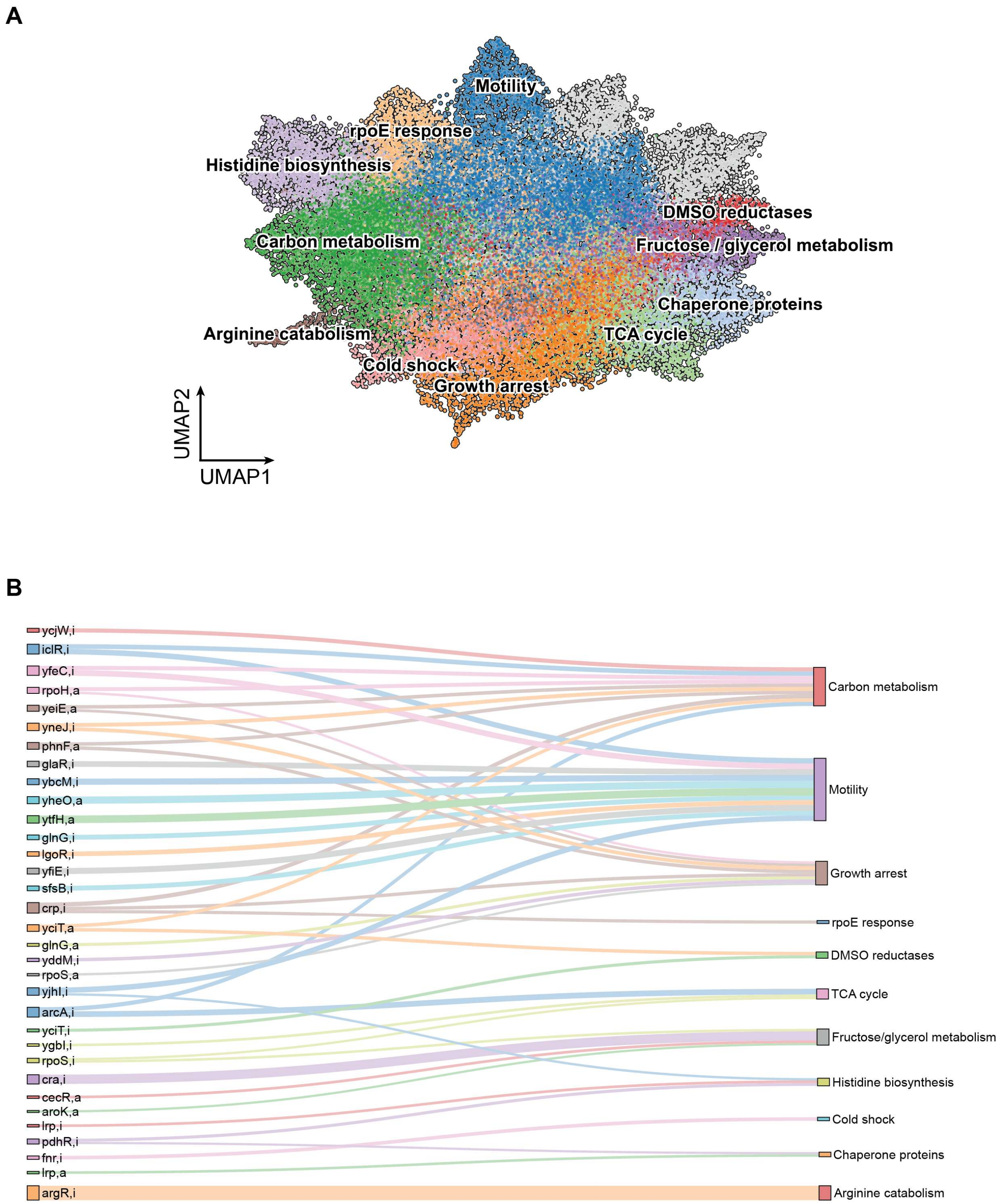


**Fig. S13. Single-cell clustering reveals convergent phenotypes across perturbations and divergence within individual perturbations. (A)** UMAP clustering where each dot represents a single cell. Clusters are colored according to Leiden designation and manually labeled using the marker gene expression profile. Clusters are colored gray if there were no perturbations overrepresented in that cluster (see Methods). **(B)** Sankey diagram highlighting the global phenotypes (UMAP clusters) exhibited by each perturbation. The stream thicknesses are proportional to the percentage of a perturbation's cells within the represented cluster.

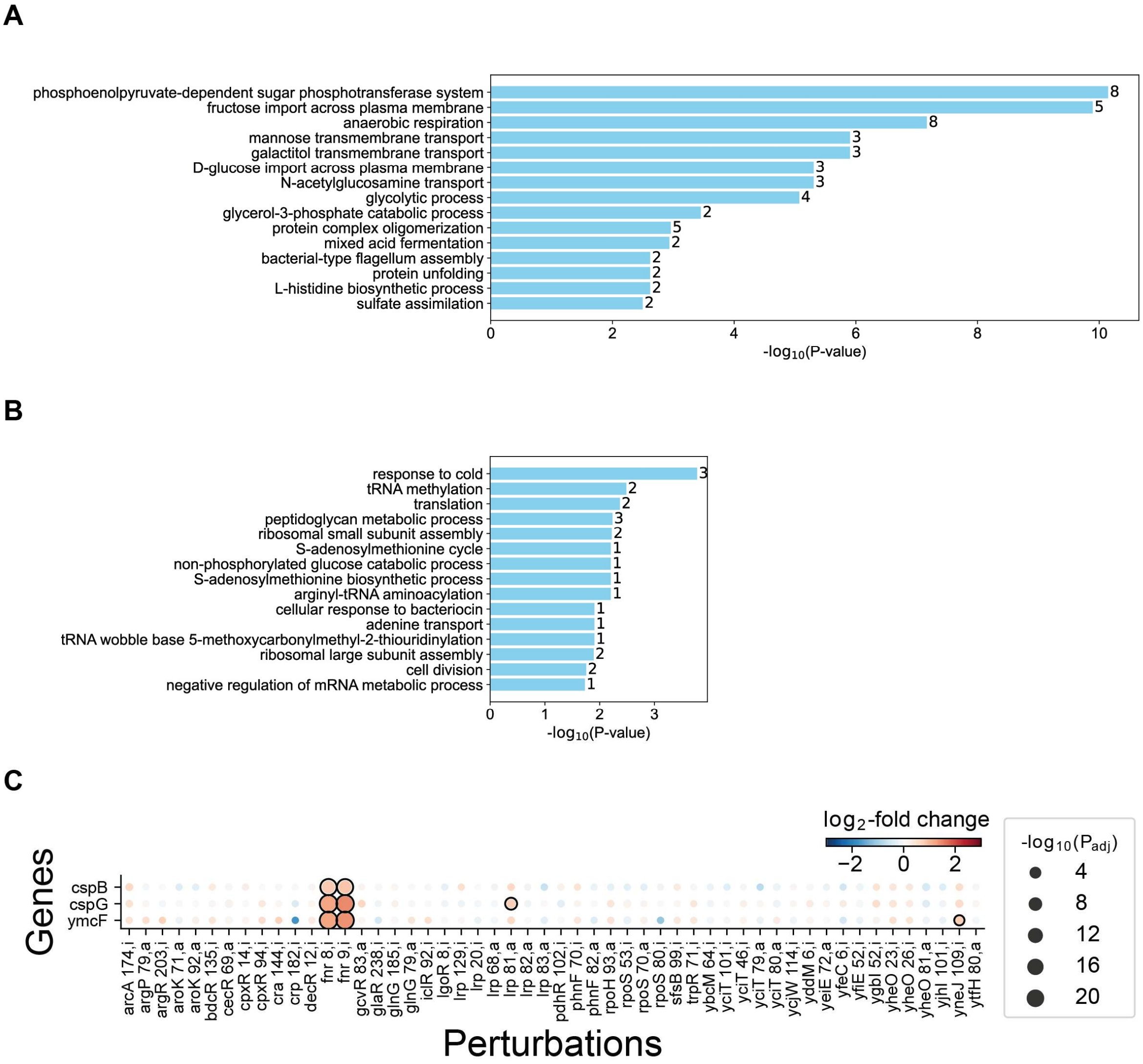


**Fig. S14. Fnr regulates cold shock response genes.** **(A)** GO-term enrichment of select biological processes calculated with the upregulated DEGs for the *fnr* CRISPRi perturbations. The p-values were calculated using a two-sided Fisher's exact test. **(B)** As in **(A)** with the downregulated DEGs. **(C)** Expression of the cold shock response genes identified in the GO-term enrichment. The dot sizes represent the adjusted p-values from the Wilcoxon rank sum test relative to an off-target control. Outlines indicate statistical significance.

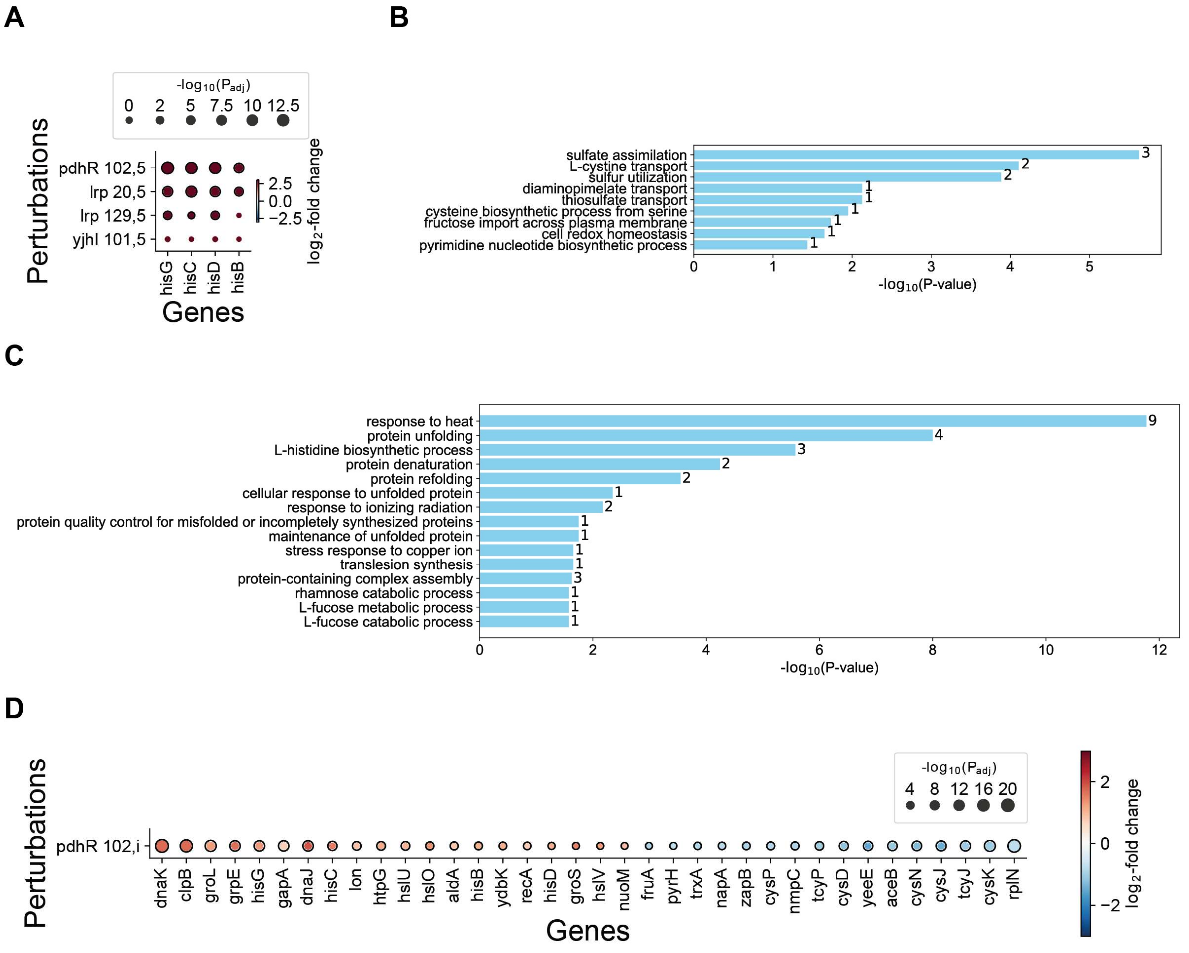


**Fig. S15. Histidine biosynthesis cluster transcriptional responses. (A)** Expression of top 4 histidine biosynthesis marker gene (ranked by the absolute z-score from a Wilcoxon rank sum test). The $\log_2$-fold changes are between the cells for each perturbation within the cluster relative to the off-target cells out of the cluster and significance for each gene is calculated from a Wilcoxon rank sum test. The dot sizes represent the adjusted p-values from the Wilcoxon rank sum test. **(B)** GO-term enrichment of select biological processes calculated with the downregulated DEGs for the *pdhR* CRISPRi perturbation. The p-values were calculated using a two-sided Fisher's exact test. **(C)** As in **(B)** with the upregulated DEGs. **(D)** Expression of CRISPRi of *pdhR* DEGs. The dot sizes represent the adjusted p-values from the Wilcoxon rank sum test relative to an off-target control. Outlines indicate statistical significance.

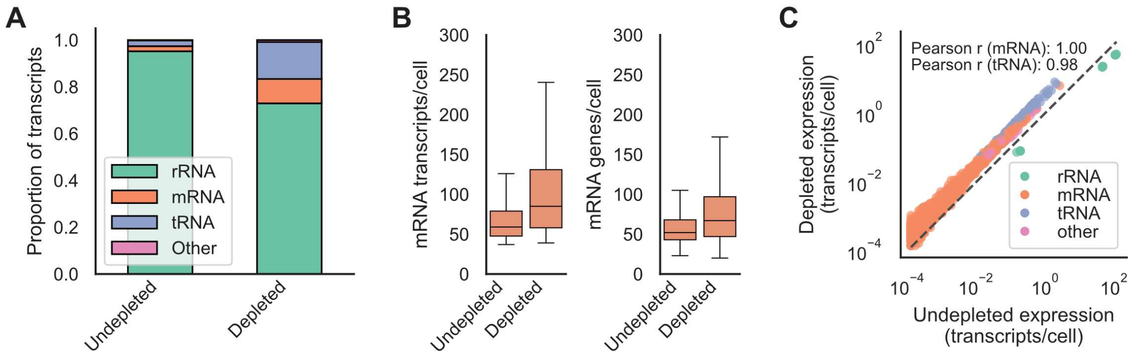


**Fig. S16. Efficacy of sequencing-guide rRNA depletion in *P. putida*. (A)** Relative proportion of RNA transcripts between samples with and without sequencing-guided rRNA depletion calculated after genome alignment but before single cell filtering. **(B)** Median and IQRs of mRNA transcripts and genes per cell with and without sequencing-guided rRNA depletion calculated after cell filtering with identical parameters. **(C)** Each unique gene was plotted with its expression in the with (y-axis) and without (x-axis) sequencing-guided rRNA depletion calculated as its number of transcripts in each library divided by the total number of filtered cells.

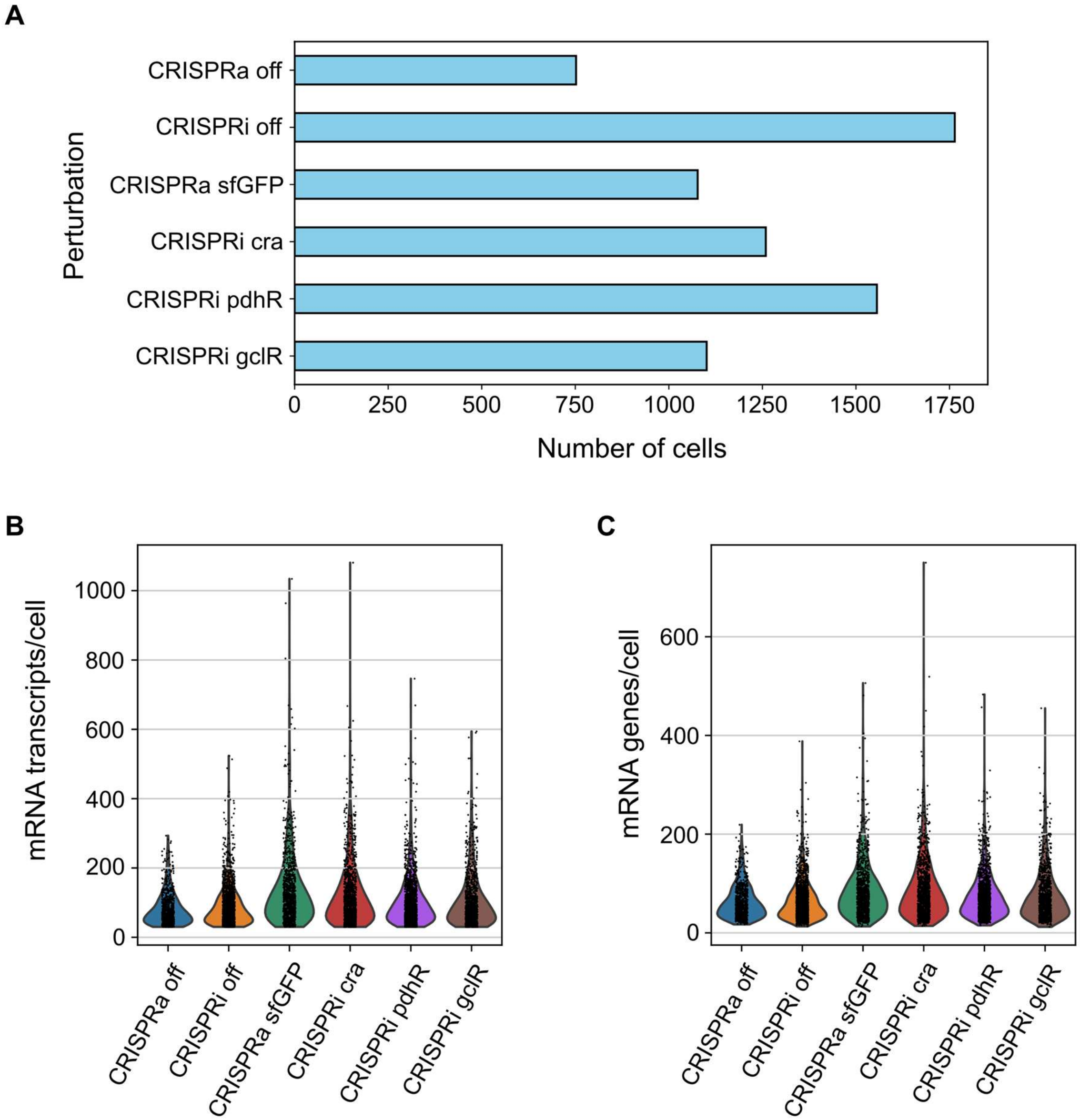


**Fig. S17. Porting mapSPLiT to *P. putida* sequencing metrics. (A)** Cell counts across CRISPR perturbations. (**B)** Distribution of transcripts per cell with each dot representing a single cell. **(C)** Distribution of genes per cell with each dot representing a single cell.
